# Linking Molecular Changes in the Hypothalamus to Alzheimer’s Disease Phenotypes

**DOI:** 10.64898/2026.11.24.768969

**Authors:** Jonathan Algoo, Catherine Kaczorowski, Amy R. Dunn, Vilas Menon, Kristen M.S. O’Connell

**Affiliations:** Center for Translational and Computational Neuroimmunology, Department of Neurology, Columbia University Irving Medical Center, New York, NY; The University of Michigan, Ann Arbor, MI; The Jackson Laboratory, Bar Harbor, ME

## Abstract

In many individuals with Alzheimer’s disease (AD), sleep disruption and weight loss emerge years before the onset of memory decline and dementia. However, the cellular and molecular bases of these early events are not well understood. The hypothalamus is a critical brain region for the regulation of these behaviors, but its response to amyloid has not been systematically characterized. We performed single-nucleus RNA sequencing to generate transcriptomic profiles of 1.1 million nuclei across the hypothalamus from 10 genetic backgrounds of the AD-BXD genetic reference panel spanning two ages and both sexes, including age-matched controls. Amyloid was associated with widespread, sex-biased molecular changes across diverse cell classes, including elevated neuroinflammatory signatures in microglia and astrocytes, upregulated ion transport pathways across barrier-associated cell types, and sex- and age-dependent changes to primary ciliary pathways in *Glp1r+/Lepr+*, Histaminergic, and *Hcrt+* neuronal populations. Two metabolism-related populations showed differential expression exclusively in one sex: *Glp1r+/Lepr+* neurons in females and *Gpr50+* neurons in males. Several findings were conserved in analysis of human hypothalamus snRNAseq data, including reduced expression of TGF-β pathway genes in medial mammillary body neurons and increased relative abundance of reactive *Serpina3n+/*YKL-40 astrocytes. This atlas provides a cellular framework for understanding the early, non-cognitive symptoms of AD and identifies several hypothalamic cell-type specific pathways that represent potential targets for therapeutic intervention.

## Introduction

The hypothalamus is a highly conserved brain region that coordinates aspects of physiology and cognition, including circadian rhythm^1^, body composition^2^, mood regulation^3^, and aspects of memory^4^. These traits are altered in Alzheimer’s Disease (AD) patients^5^, which suggests a link between AD and dysfunctional cellular physiology in the region, though this hypothesis has largely been untested. While prior large-scale molecular profiling studies have identified cell type-specific molecular signatures of AD in the cortex^6,7^ and hippocampus^8^, there have been few studies that have used modern molecular methods such as single-cell (scRNAseq) or single-nucleus RNA sequencing (snRNAseq) to profile the hypothalamus using tissue from human AD patients or animal models of AD.

Accumulating clinical and molecular evidence motivates a more comprehensive cellular investigation. At a structural level, hypothalamic atrophy is progressive throughout AD pathogenesis, and volume loss correlates with cognitive decline and sleep alterations^9^. Neuropeptides, key signaling molecules that coordinate hypothalamic function, exhibit different concentrations in the cerebrospinal fluid (CSF) of AD patients^10,11^. Weight loss^12,13^ and sleep cycle dysregulation^9,14^ often precede cognitive decline and manifest before an official AD diagnosis.

A thorough investigation of the molecular content of the hypothalamus must overcome several technical challenges. The region is highly heterogeneous and is composed of approximately a dozen nuclei (subregions)^15^; several contain rare, but physiologically important, neuronal populations exhibiting distinct neuropeptide and receptor profiles^16^. Attaining a comprehensive assessment of these types is important as neuropeptide networks coordinate physiology both within and between nuclei; observational human and experimental animal model studies have described how dysfunctional peptide signaling leads to systemic effects on homeostasis^17^. Profiling the hypothalamus is further complicated as some cell types exhibit low expression of biologically important neuropeptides or receptors^16^. To profile these genes, sn/scRNAseq studies must employ high sequencing depths per cell, which is challenging when coupled with the need to profile large numbers of cells to ensure representation of rare populations. These constraints are reflected in prior work as the only two human^18,19^ studies of the hypothalamus in AD have either selected a single subregion (the mammillary bodies) with a larger number of samples or used relatively modest sample sizes across multiple subregions. Similarly, the small number of animal studies that have applied modern molecular omics methods to study the hypothalamus with AD models^20–22^ have profiled few samples from single, inbred genetic backgrounds, leaving contributions of the region to AD risk largely uncharacterized.

Animal models are particularly well-suited to investigating the role of the hypothalamus in AD. In contrast to post-mortem human studies, they enable larger sample sizes, the ability to assess molecular changes at relatively early timepoints of disease progression, and environmental control. The latter is especially valuable when investigating traits like weight and sleep disruption, which are heavily influenced by environmental factors that complicate efforts to isolate the relationship between AD and these traits in observational human studies. However, a downside of using conventional animal models is that they often fail to translate to human biology, due in part to the lack of genetic diversity in common models of AD. Given the strong effects of genetic composition on genotype-phenotype interactions^23^, findings from studies that use single-strain models are restricted to a specific genetic background and thus have limited generalizability to human populations. In this study, we explicitly addressed this deficit by profiling mice from the AD-BXD panel^24^ of genetic diversity. Our findings thus capture generalizable disease-associated signals, enabling a more representative investigation of the cellular and molecular effects of amyloid.

To investigate the effects of amyloid on the hypothalamus, we generated an snRNAseq resource composed of 1.1 million high-quality and high-read depth nuclei across the entire hypothalamus from 79 mice, balanced across age (6 and 14 months), sex, AD transgene status (5XFAD or non-transgenic/Ntg), and 10 genetic backgrounds. This dataset constitutes the first large-scale molecular profile of the hypothalamus in a genetically diverse preclinical model of AD, and represents a substantial addition to the total number of hypothalamic cells profiled via sc/snRNAseq to date.

Comparing transgenic to control animals, we identify non-neuronal signatures common to other regions, such as higher levels of activated microglia and reactive astrocytes, as well as new AD associations including altered ion transport expression across cell types forming the brain-CSF, blood-brain, blood-CSF, and brain-meninges barriers. We additionally characterize changes to neuronal populations with established roles in sleep, metabolism, and memory, and identify multiple neuronal types exhibiting sex- and age-dependent expression changes of primary cilia assembly. Several findings were validated using human snRNAseq data, including dysregulated genes in microglia, downregulated levels of TGF-β genes in medial mammillary body neurons, and higher levels of an inflammatory astrocyte state marked by *CHI3L1* and *SERPINA3*. Ultimately, this investigation delineates plausible cellular and molecular bases for several cognitive and non-cognitive symptoms observed in AD.

## Results

### Construction and Description of a Hypothalamic Single Nucleus Atlas

We performed snRNAseq on hypothalami isolated from a cohort of 79 mice, evenly balanced across age, sex, AD transgene status, and 10 genetic backgrounds (Fig. 1A, 1B). The AD-BXD and control animals profiled here are a subset of a larger population of mice that have been assessed for behavioral and physiological measurements throughout the lifespan^25^. These mice, alongside other animals matching the genetic backgrounds profiled in the present study, present changes to cognitive (contextual fear acquisition/CFA assay performance) and homeostatic (weight and sleep) traits in response to the 5XFAD transgene (Fig. 1C) that mirror the human AD phenotypes of decreased memory performance, weight, and sleep time.

**Figure 1.**
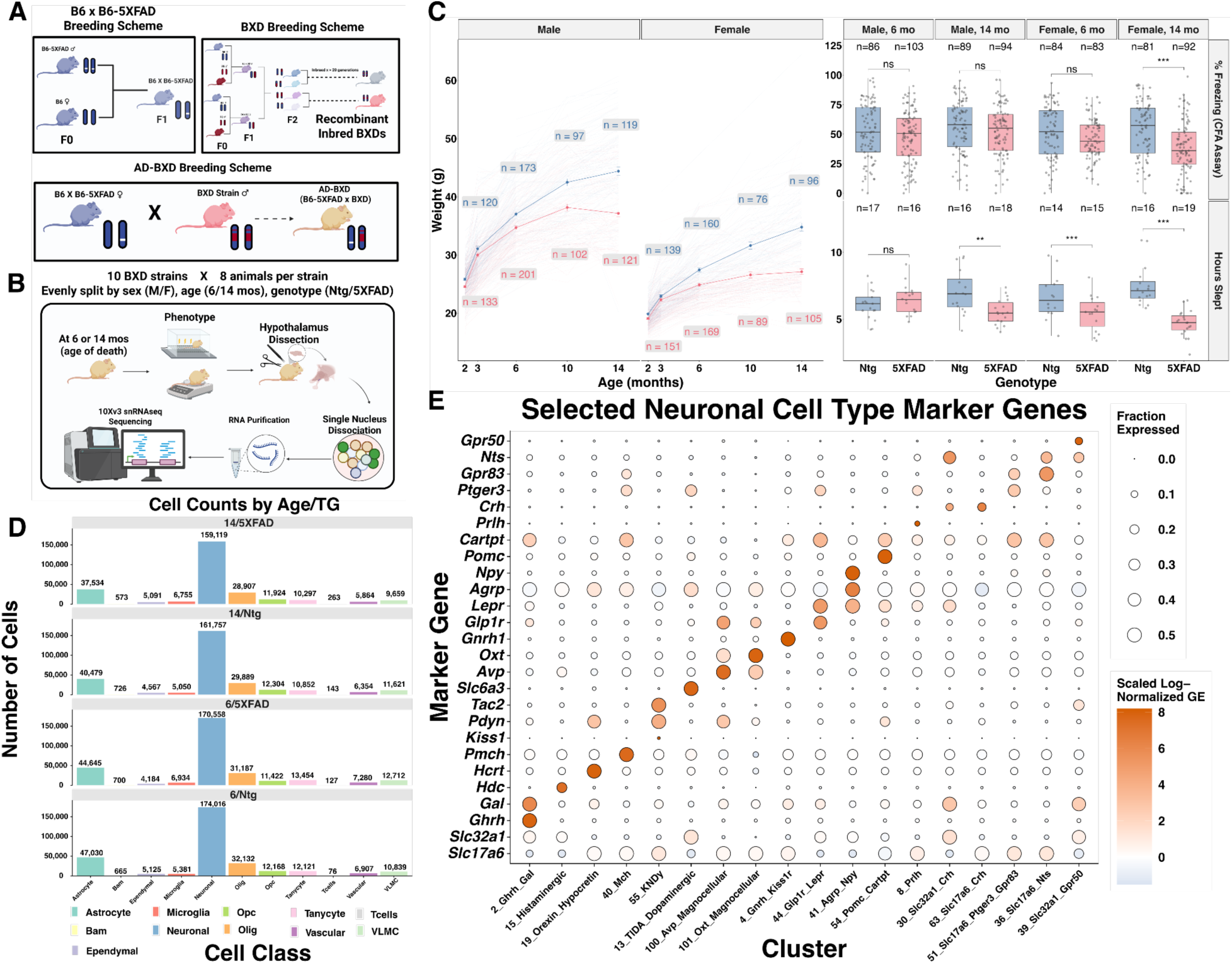
Generation of a single nucleus atlas of the hypothalamus across age (6/14 months), sex, transgene status (5XFAD/Ntg), and 10 genetic backgrounds. A) Overview of the AD-BXD breeding scheme. B) Experimental workflow. Mice were assessed for cognitive and homeostatic phenotypes, including contextual fear acquisition and body weight. Bilateral whole hypothalami were extracted and profiled with snRNAseq. C) The presence of the 5XFAD transgene (red line/boxplot) significantly influences CFA performance, body weight, and total sleep time, mirroring human AD phenotypes. Statistical significance for CFA performance and sleep time was assessed for each age/sex separately using a linear model. P-values were adjusted across all comparisons tested (grouping across both phenotypes, age, and sex) through Benjamini-Hochberg FDR^27^ (FDR) correction. Non-independence of observations (due to strain) was accounted for via a clustered covariance matrix with the HC2 estimation type. D) Bar plots of the number of nuclei assigned to each cell type within each age/transgene group. Similar numbers per cell type were present across groups. E) Multiple characterized neuronal subtypes are identifiable by marker gene expression. Color indicates z-scored log-normalized gene expression, and size of dot indicates fraction of cells with non-zero expression (fraction expressed capped at 0.5 for visualization). Opc = Oligodendrocyte precursor cell; Bam = Border Associated Macrophages; Olig = Oligodendrocyte; VLMC = Vascular and Leptomeningeal Cell. Panels A and B created with Biorender.

After quality control and clustering (Methods), we produced a single-nucleus atlas comprising 1.1 million nuclei across 41 glial and 69 neuronal clusters. We observed a median of 8997 unique molecular identifiers (UMIs) and 3553 genes detected across all nuclei which reflects the high quality of this dataset. There was substantial variability in these metrics between cell classes (Supplementary Fig. 1A,B), which was expected and consistent with other transcriptomic studies of the brain^26^. Across cell types, we observed large numbers of cells across all four age x transgene groups (Fig. 1D). Overall, this dataset contains at least 2-3 fold more nuclei profiled than other relevant studies of the mouse hypothalamus (Supplementary Fig. 1C).

As expected, neuronal clusters were partitioned into GABAergic or glutamatergic populations, with one cluster expressing markers for both neurotransmitters. Some neuronal clusters corresponded to established neuropeptide-expressing cell types (Fig. 1E), although the neuronal populations that have been comprehensively investigated for their role in physiological functions are a minority of all clusters observed (Supplementary Fig. 2). We characterized extensive heterogeneity in neuropeptide expression, including within established cell types, which highlights the interconnected nature of hypothalamic peptide signaling and the potential pleiotropic effects of neuropeptides to impact multiple aspects of physiology. We additionally identified large numbers of glia located within the parenchyma (oligodendrocytes, astrocytes, oligodendrocyte precursor cells (OPCs), microglia, vascular cells) and barrier cell types that form or enforce borders within or surrounding the hypothalamus (border associated macrophages (BAMs), *glia limitans superficialis* (GLS) astrocytes, vascular and leptomeningeal cells (VLMCs), ependymal cells, and tanycytes).

Most preclinical studies have focused on one or a few inbred strains and there has been little investigation into the effects of genetic heterogeneity on the molecular and cellular landscape of the hypothalamus; therefore, we first evaluated the impact of genetic diversity on the molecular state of the hypothalamus. We were particularly interested in whether genetic background influences cell type-level variability, as both hypothalamic function and AD pathophysiology are driven by cell-type specific mechanisms. We performed a variance partitioning analysis to evaluate the association between strain and gene expression, identifying 612 genes across cell types where strain contributed 30% or more to total variance (Fig. 2A, Supplementary Table 1). Read mapping percentages were not different across strains, suggesting this result is not a technical artifact due to differential alignment to the B6 reference genome across strains (Supplementary Fig. 1D). There was variability in the number of cell type-specific strain-associated genes: GABAergic neurons exhibited the largest number of strain-variable genes, while vascular cells exhibited the lowest (Supplementary Table 2). Most genes exhibited variability in only one cell type, although 60 were shared across 3 or more types (Supplementary Table 1); notably, one variably expressed gene was the receptor for glucagon-like peptide 1 receptor (*Glp1r*), best known as the target for semaglutide and other Glp1r agonists (Supplementary Table 1). We further evaluated whether genetic background is associated with cellular composition and found several robust differences across cell types between the most commonly used laboratory strain, C57BL/6J (B6), and the BXD F1 strains after accounting for age, sex, and genotype (Supplementary Fig. 3). Overall, the key result from this analysis is the identification of cell-type level molecular heterogeneity in the hypothalamus across genetically distinct mouse strains.

**Figure 2.**
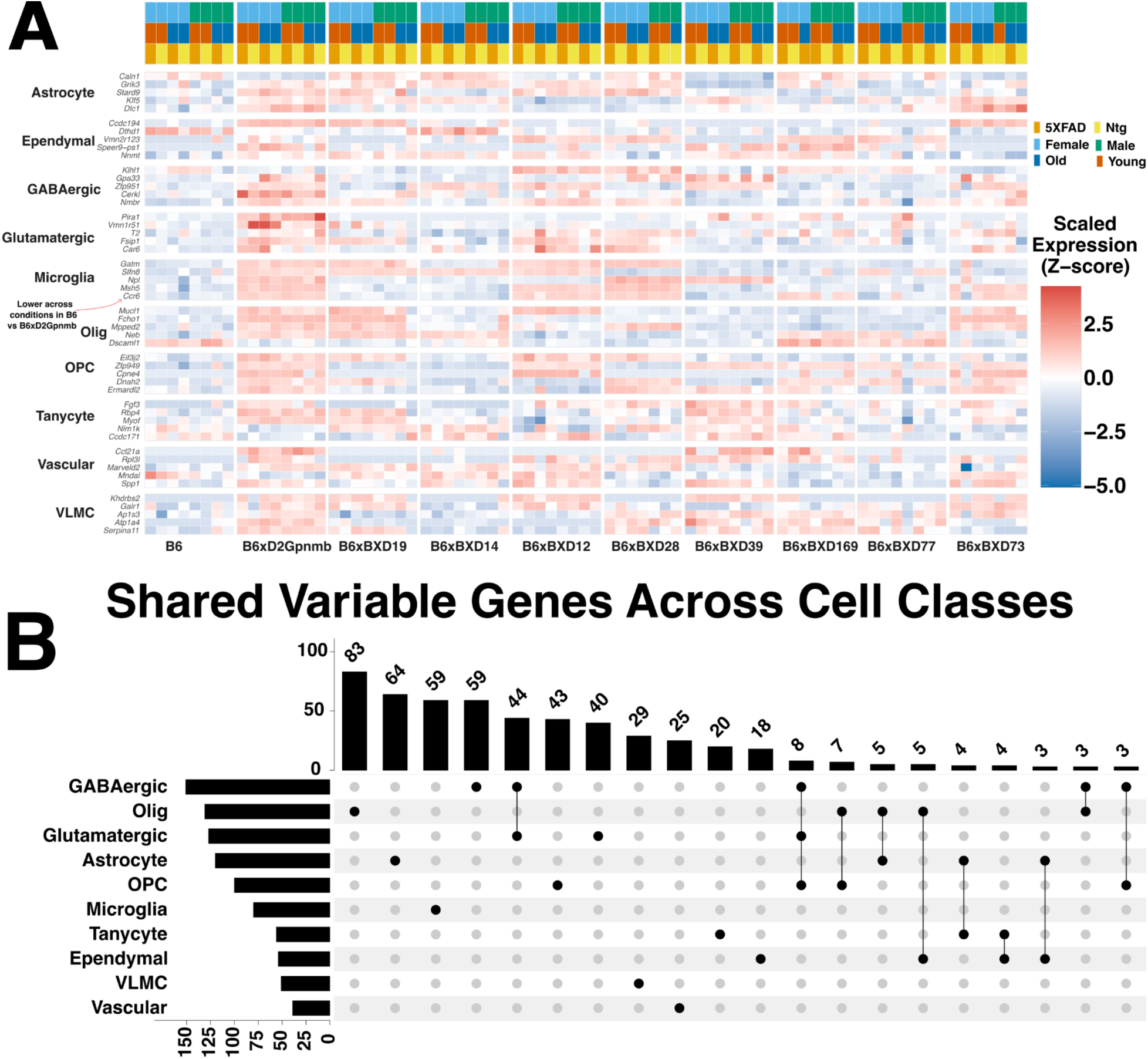
Genetic background leads to non-negligible molecular heterogeneity at the cell type level. A) Cell- type specific variable gene expression, with average scaled expression (across all cells of a given class) visualized for each cell class (y-axis) and strain (x-axis). Expression of some physiologically important genes, such as Ccr6 in microglia, is strongly influenced by genetic background across all age/sex/transgene status groups. B) Upset plot displaying number of variable genes corresponding to individual cell types. GABAergic neurons exhibit the highest number of strain-regulated variable genes, followed by oligodendrocytes, glutamatergic neurons, and astrocytes. Some variable genes are shared across cell types, such as the 44 common to both glutamatergic and GABAergic neurons. OPC = Oligodendrocyte precursor cell; Olig = Oligodendrocyte; VLMC = Vascular and Leptomeningeal Cell.

### Increased Relative Abundance of Disease Associated Subtypes and Levels of a Sex-Biased MHC Signature in Hypothalamic Microglia

Microglia are the primary immune cell within the central nervous system and are widely recognized for strong disease-modifying effects that contribute to AD pathogenesis. Microglia were assigned to defined subtypes based on expression of cluster-specific, inflammatory (*P2ry12*, *Tmem119*, *Cx3cr1*, *Trem2*, *Apoe*), and sex-specific (*Xist*, *Tsix*, *Uty*) marker genes (Fig. 3A). Two clusters, microglia-4 and microglia-5, exhibited signatures of disease-associated microglia (DAMs). DAMs have been hypothesized as conserved sensors of neurodegeneration^28^ and have been identified in other brain regions in the 5XFAD model^29^. Microglia transition from a resting state into DAMs via a two-stage process (homeostatic to Stage 1 DAMs, and Stage 1 to Stage 2 DAMs); Stage 1 and Stage 2 DAMs exhibit distinct transcriptional signatures. Microglia-4 exhibited higher expression of previously defined Stage 2 DAM markers^28^ whereas microglia-5 better corresponded to Stage 1 DAMs (Supplementary Fig. 4A).

**Figure 3.**
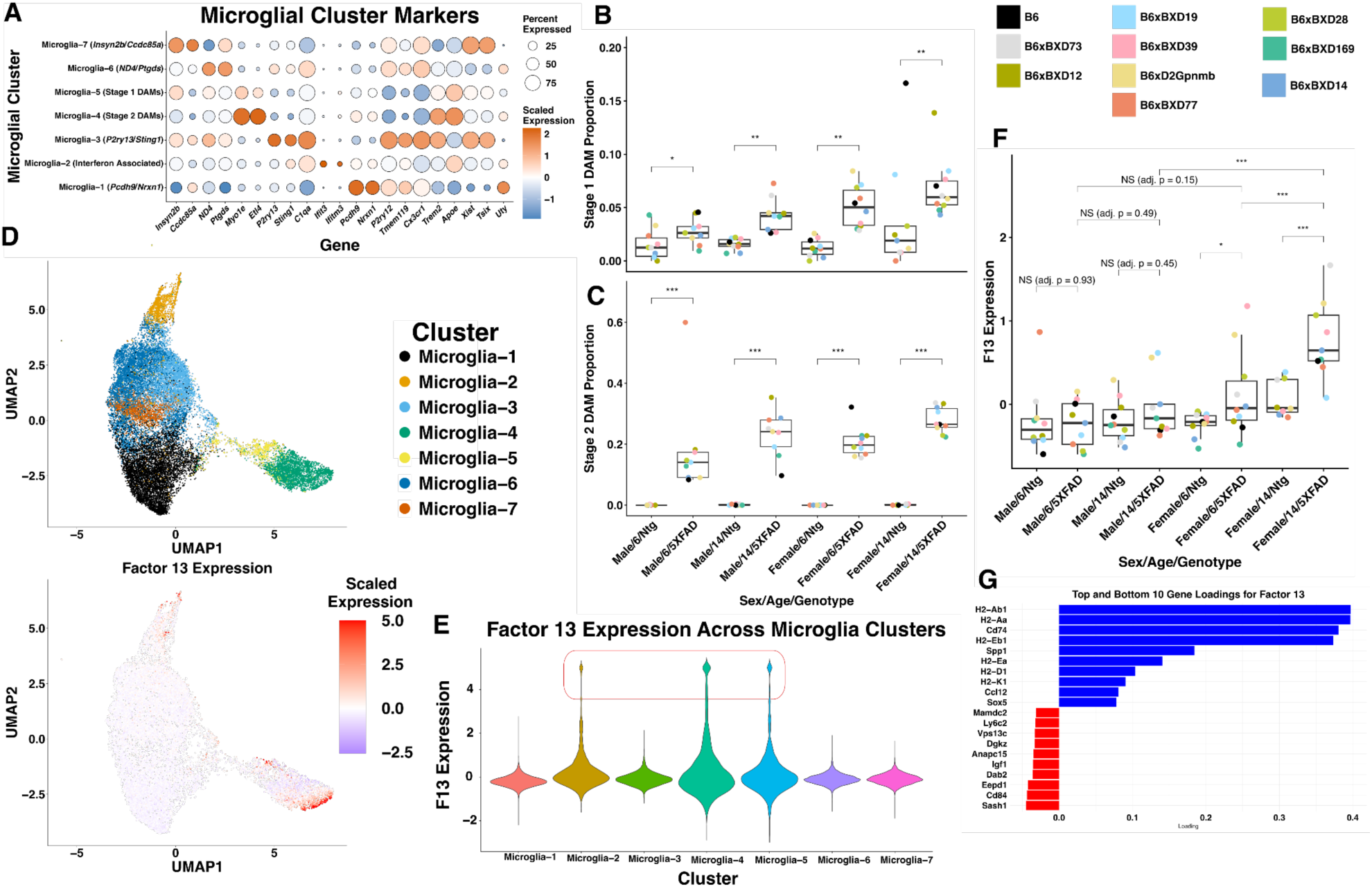
Microglia exhibit broad transgene-dependent transcriptomic shifts and cluster-spanning pathway level changes. A) Dotplot of cluster-distinguishing marker genes (left) and microglial activation, AD-associated, and sex-biased marker genes (right). B,C) Stage 1 (microglia-5) and Stage 2 (microglia-4) DAMs are found at higher levels in 5XFAD mice of both sexes. D,E) Factor 13 exhibits high expression in subsets of microglia-2, microglia-4, and microglia-5. Expression overlaid on UMAP coordinates (D) and via a violin plot across clusters (E). Factor 13 expression was clipped between 5 and −5 for the UMAP and violin plot. F) Factor 13 (F13) exhibits sex, age, and transgene-dependent expression changes. This visualization and analysis only included aggregate factor expression from animals with at least 50 microglial cells profiled. G) Factor 13 exhibits high loadings of several MHC genes, like H2-Ab1 and H2-Aa.

We evaluated whether the 5XFAD transgene caused changes to the relative abundance of microglial clusters. Given that individual differential composition methods exhibit variable results when applied to the same datasets^30,31^, we used a consensus approach combining three separate methods to ensure robustness (Methods). The 5XFAD transgene was associated with an increased relative abundance of microglia-4 and microglia-5 in both sexes in both 6 and 14 month mice (Fig. 3B, 3C; Supplementary Table 3). As an orthogonal replication, we verified upregulated levels of Stage 1 and 2 DAMs, mapping microglial cluster assignments onto a previously published MERFISH dataset that profiled 12 month 5XFAD and Ntg mice with a B6 genetic background, alongside hemizygous 5XFAD/heterozygous *Trem2*^R47H^ and Ntg/heterozygous *Trem2*^R47H^mice of both sexes^22^ (Supplementary Fig. 5A,B; Methods).The authors that generated the MERFISH dataset previously noted a higher relative abundance of DAMs broadly within the hypothalamus alongside other brain regions; our analysis generalizes this finding to genetically diverse models and specifically characterizes higher levels of both Stage 1 and Stage 2 DAMs within the hypothalamus.^22^

Enforcing discrete cluster assignments can limit the detection of transcriptional shifts manifesting across supra-cluster resolutions. This restriction is particularly acute for dynamic cell types like microglia, where environmental stimuli can result in continuous transcriptional gradients. To complement our cluster-based analysis, we also performed de novo factor analysis^32^ to discover correlated patterns of population-wide transcriptomic variation independent of cluster boundaries and identified 50 dimensions of variation (Supplementary Tables 4,5). Some factors, like Factor 2 (upregulated in the interferon associated subtype), corresponded to individual clusters, but many exhibited more complex patterns of variation across discrete transcriptomic clusters (Supplementary Fig. 4B).

We evaluated associations between each factor and transgene status, identifying 73 statistically significant amyloid-related associations across all comparisons tested (Supplementary Table 6). Notably, Factor 13 - which exhibited high loadings for the immunomodulatory gene *Spp1* (the gene encoding for osteopontin) and several H2 genes of the major histocompatibility complex (MHC) such as *H2-Ab1*, *H2-Aa*, *H2-Ea*, *H2-D1*, and *H2-K1* (Fig. 3G) - demonstrated sex-dependent associations with amyloid pathology. Female 5XFAD mice showed upregulation of Factor 13 compared to their non-transgenic counterparts at both 6 months (adj. p < 0.05) and 14 months (adj. p < 0.001), whereas male 5XFAD mice exhibited no significant changes compared to their non-transgenic counterparts (6 month adj. p = 0.93; 14 month adj. p = 0.45). A direct comparison revealed higher levels in older transgenic females than males (adj. p < 0.001), an association that was not significant at 6 months (adj. p = 0.15), indicating that altered levels of this pathway in females are influenced by age (Fig. 3F). We next investigated which clusters are enriched for Factor 13; high levels of Factor 13 are restricted to subpopulations of interferon-associated microglia and DAM clusters (Fig. 3D, Fig. 3E). Together, these results identify a strong sex-specific MHC response that manifests across discrete microglial clusters.

### Chi3l1+/Serpina3n+ Astrocyte Cluster Relative Abundance is Higher in 5XFAD mice

Astrocytes can transition to a reactive state via inflammatory compounds secreted by activated microglia, and the presence of activated microglia has been identified as a mediator of astrocytic reactivity in human AD.^33^ Given this intercellular crosstalk, and the identification of higher microglial activation signatures, we hypothesized astrocytes would also display an upregulated inflammatory response. We characterized 9 transcriptionally distinct clusters that exhibit specific marker gene profiles, and which additionally segregated by expression of reactivity-associated and immediate early genes (Fig. 4A). Three clusters (astrocyte-2, astrocyte-3, astrocyte-5) exhibited reactivity signatures, while astrocyte-5, which was marked by high levels of the AD-associated genes *Serpina3n* and *Chi3l1*, was additionally more abundant in 5XFAD animals of both sexes at 6 and 14 months (Fig. 4B; Supplementary Table 3).

**Figure 4.**
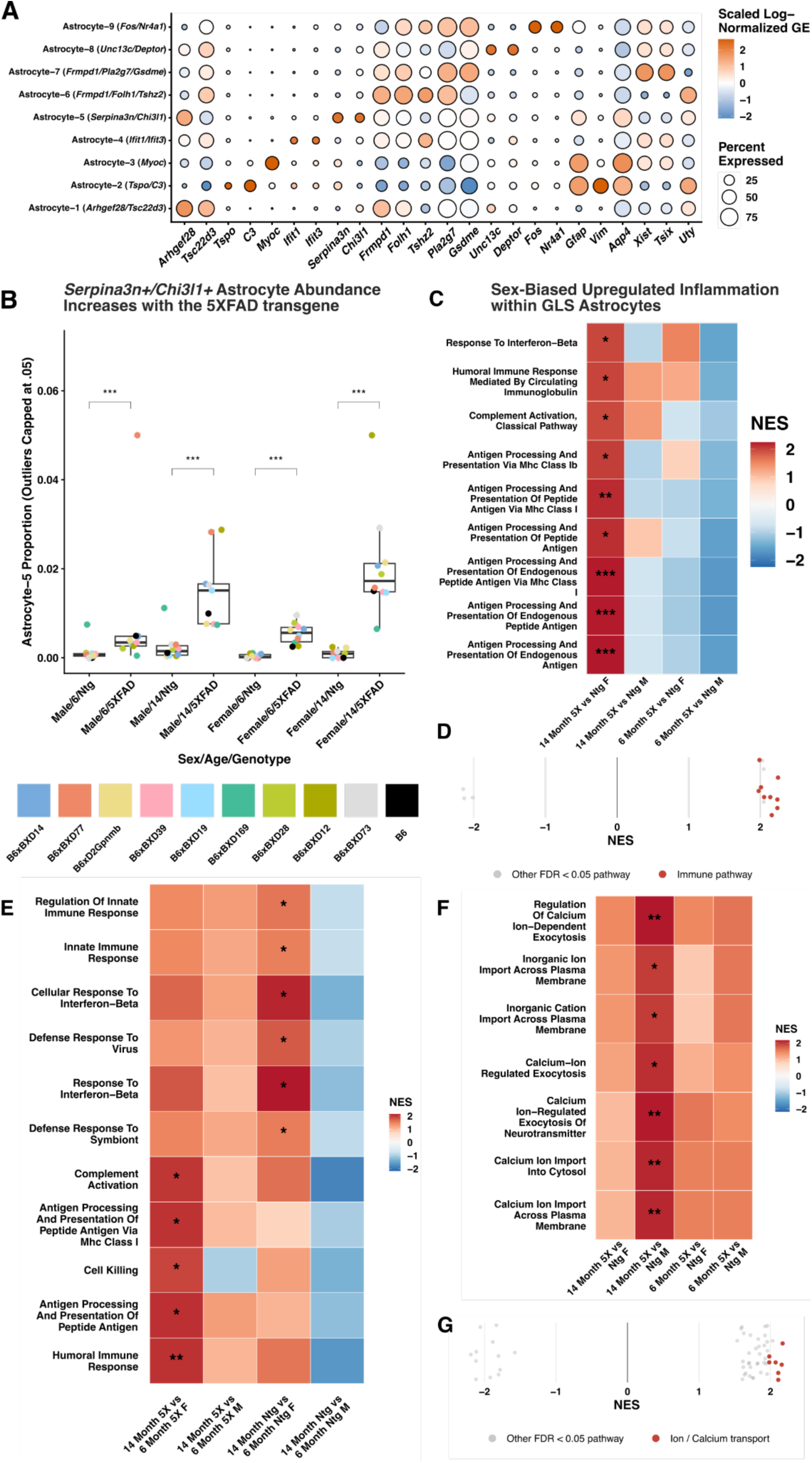
Astrocytes exhibit ion transport expression and sex-biased inflammatory associations with the 5XFAD transgene. A) Cluster-distinguishing marker genes (left) alongside canonical astrocyte and sex-biased marker genes (right). B) Relative abundance of astrocyte-5 (Serpina3n+/Chi3l1+) is higher in transgenic 5XFAD animals in both males and females at 6 and 14 months. Max value set at 0.05 for visualization. C) Upregulated GLS immune signatures are detectable only in comparisons of transgenic versus nontransgenic animals in 14 month females. D) Some of the strongest upregulated signatures in GLS astrocytes between 14 month 5XFAD and Ntg female animals are related to immunity. E) GLS astrocytes exhibit sex-biased immune signatures in aging, with and without the 5XFAD transgene. F) Several ion transport pathways are strongly upregulated in male 14 month 5XFAD animals versus nontransgenic controls and G) exhibit large normalized enrichment scores (NES) relative to other statistically significant pathways.

### GLS Astrocytes Exhibit Sex-Dependent Inflammatory and Ion Transport Signatures

A specialized population of astrocytes known as *glia limitans superficialis* (GLS) form a barrier that separates the parenchyma and meninges and are thought to protect the brain from peripheral pathogens and other insults^34^. The GLS population is not well understood, and to our knowledge no prior studies have directly characterized changes to this population using sc/snRNAseq in human AD or an AD animal model. In our data, cells from the astrocyte-3 cluster are enriched for *Myoc* (Fig. 4A), a marker gene that uniquely marks GLS astrocytes brain-wide^34^. Given the expected lower degree of heterogeneity within an individual cluster, we focused differential expression analysis at the individual gene level (Supplementary Table 7). We then performed gene set enrichment analysis (GSEA)^35^, a technique that assesses whether predefined sets of genes in a pathway are selectively enriched at either end of a list of genes ranked by fold change. In females, most of the pathways that were upregulated in GLS astrocytes in a comparison of 14 month 5XFAD animals vs Ntg controls were related to inflammation (Fig. 4C and D; Supplementary Table 8); however, GLS astrocytes from males did not exhibit significant inflammatory associations at either 6 or 14 months and these signatures were not detectable in females at 6 months (Fig. 4C). The significant signatures may reflect exacerbated processes of female midlife aging as several inflammatory pathways were upregulated in comparisons of 14 month to 6 month animals both with and without the 5XFAD transgene (Fig. 4E).

In comparisons of male 5XFAD vs Ntg animals at 14 months, several statistically significant pathways with the largest effect sizes were related to ion transport (Fig. 4F and G). These signatures were only significant in male 14 month 5XFAD vs Ntg animals and not 6 month male 5XFAD vs 6 month Ntg or any female comparisons, although they exhibited similar directional shifts (Fig. 4F). Multiple pathways specifically implicated calcium transport. In prior work in a related cell type, astrocytes of the blood brain barrier/*glia limitans perivascularis*, calcium dynamics have been linked to endothelial cell diameter^36^. Combined with the potent role of calcium dynamics broadly in regulating astrocytic activity^37^, we speculate dysregulated calcium activity in GLS astrocytes may reflect altered function.

### Expression of Ion Transport Pathways is Upregulated Due to the 5XFAD Transgene Across Barriers and Cell Types

Motivated by the discovery of ion transport expression changes in GLS astrocytes, we investigated whether other cell types that form or regulate the different barriers of the hypothalamus (the blood/brain, brain/meninges, blood/CSF, and brain/CSF barriers) exhibited similar changes. In all barriers, we identified cell types with higher ion transport expression in at least one sex in 14 month comparisons; several cell types exhibited significant changes to the same pathways at 6 months (Fig. 5 A-D; Supplementary Fig. 7; Supplementary Tables 9,10). In several cell types (such as two subtypes of ependymal cells), ion transport pathways were among the most upregulated based on normalized enrichment score (NES) (Bottom of Figs. 5 A-D).

**Figure 5.**
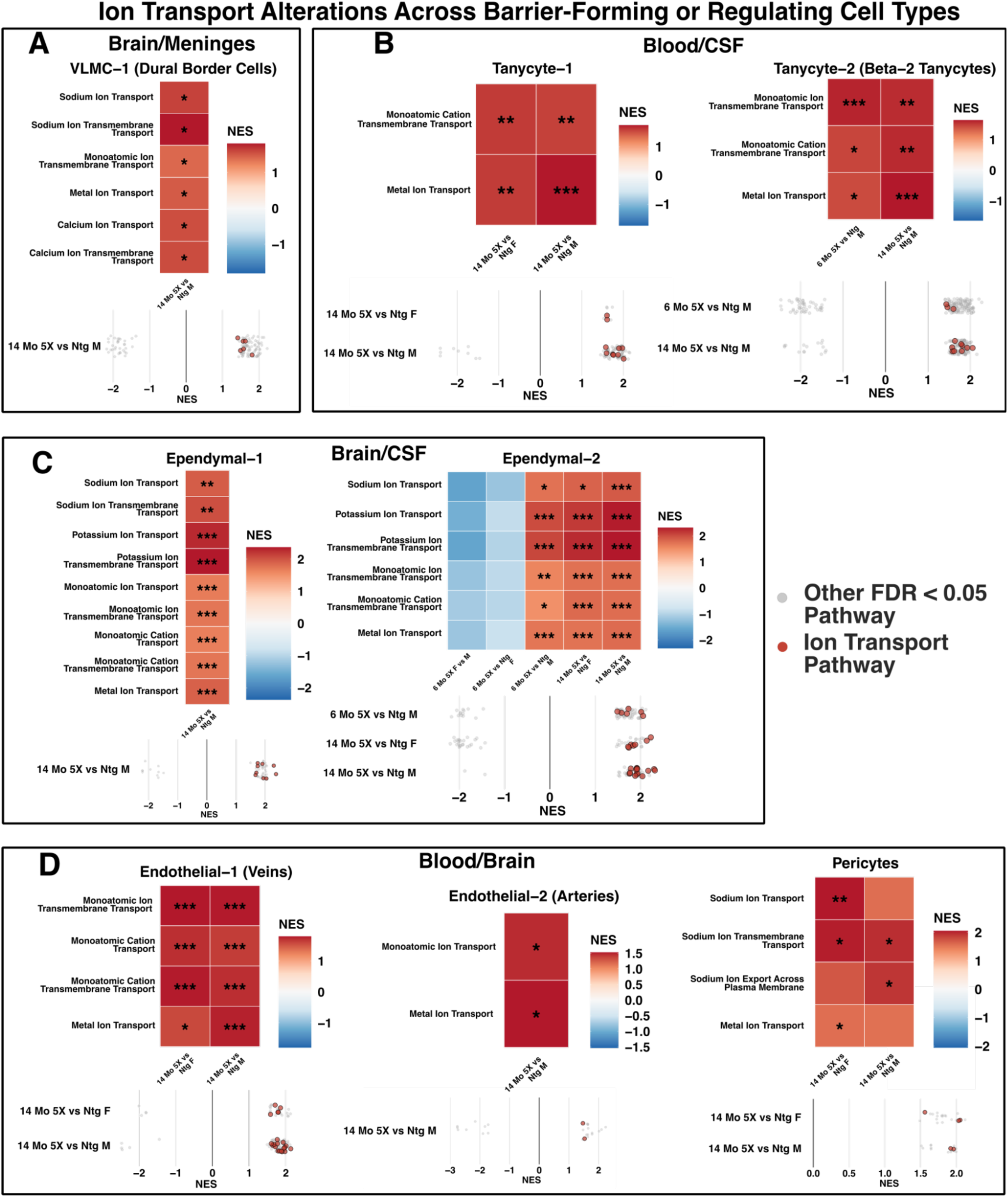
Ion transport pathway expression is upregulated due to the 5XFAD transgene across multiple barriers in the hypothalamus. The 5XFAD transgene is associated with upregulation of several ion transport pathways, including A) sodium, metal, and calcium ion transport pathways in dural border cells of the brain/meninges barrier; B) cation and metal ion transport pathways in two tanycyte subtypes (including beta-2 tanycytes of the median eminence); C) potassium ion and metal ion transport pathways in two ependymal cell subtypes; D) metal ion transport pathways in venous and arterial endothelial cells, and sodium ion transport pathways in pericytes of the blood-brain barrier. For each cluster, the top heatmap displays selected significant ion transport pathways, and the bottom strip plot highlights the NES scores of all altered ion transport pathways (red dots), including those that are not highlighted in the top panel, relative to other significant pathways (gray dots).

At the brain/meninges barrier, VLMC-1 (dural border cells, Supplementary Fig. 6A) exhibited increased sodium, metal, and calcium ion transport pathways in 14 month male 5XFAD animals versus Ntg animals (Fig. 5A).

Ependymal cells and tanycytes line the ventricles and separate the CSF from the parenchyma; these cells were captured in our dissection as the hypothalamus surrounds the walls of the third ventricle. Different tanycyte subtypes regulate and facilitate transfer of bioactive compounds from blood to the CSF, and CSF to the parenchyma.^38^ Of the three tanycyte clusters (Supplementary Fig. 6B) in this dataset, two (tanycyte-1 and tanycyte-2) exhibited ion transport upregulation due to the 5XFAD transgene at 14 months. Changes to metal ion and cation transmembrane transport expression were observed in tanycyte-1 in 5XFAD vs Ntg 14 month comparisons of both sexes. Elevated metal ion/cation transmembrane transport expression was also observed in 6 and 14 month 5XFAD vs Ntg comparisons in males in tanycyte-2 (Fig. 5B). Tanycyte-2 expresses marker genes (specifically *Col25a1*/*Fndc3c1*)^39^ that correspond to beta-2 tanycytes, an important subpopulation located at the base of the third ventricle that facilitates sensing and transmission of molecules from blood to the CSF.^38^ The other two clusters of tanycytes, tanycyte-1 and tanycyte-3, exhibit overlapping markers of previously reported subtypes^40^ that line the lateral walls of the third ventricle and contact the parenchyma (Supplementary Fig. 6B).^39^

We identified three ependymal clusters (Supplementary Fig. 6C), two of which exhibited ion transport alterations between 14 month 5XFAD and Ntg animals of either sex (Fig. 5C). Both of these ependymal clusters exhibited sodium, potassium, and metal ion transport changes; however, ependymal-1 associations were significant only in males at 14 months, while several ependymal-2 associations were significant in both sexes at 14 months and in males at 6 months. This suggests males exhibit transport alterations earlier in ependymal-2 cells.

Finally, we identified two subtypes of endothelial cells, endothelial-1 (veins) and endothelial-2 (arteries), associated with the blood-brain barrier (Supplementary Fig. 6D). Endothelial-1 exhibited upregulated cation and metal ion transport pathways in both sexes at 14 months (Fig. 5D), while endothelial-2 ion transport pathways (monoatomic and metal ion transport) were only upregulated in males at 14 months. Additionally, pericytes, which contribute to enforcement and regulation of the blood brain barrier, exhibited elevated sodium ion transport pathway expression in both sexes at 14 months. Collectively, these pathway signatures indicate widespread changes to ion transport in several interface-forming cell types due to the 5XFAD transgene, and indicate some barrier cell subtypes, such as ependymal-2, exhibit ion transport expression changes in a sex and time-dependent manner.

### The 5XFAD Transgene Dysregulates Neuronal Types that Influence Memory, Circadian Rhythms, and Energy Balance

Changes to hypothalamic microglia, astrocytes, and barrier cell types such as tanycytes and endothelial cells have been linked to metabolic, cognitive, or physiological dyshomeostasis through their interactions with neurons^41^. We hypothesized neuronal clusters would also exhibit transcriptional changes in transgenic animals. We performed pseudobulk DEG analysis across all cell types, grouping glia at the class level and neurons at the cluster level (Supplementary Table 7). A minority of statistically significant genes with altered expression are found within 5 Mb of the 5XFAD transgene insertion location on chromosome 3 near 63 Mb^42^, and therefore may be technical artifacts (ex: the *Vmn2r1* gene was dysregulated in 28 cell types in males and 9 in females in 14 month 5XFAD vs Ntg comparisons). Counting the numbers of genes with altered expression patterns across clusters (excluding genes within 5 Mb of the transgene), microglia exhibited the highest numbers of DEGs at both 6 and 14 months (Fig. 6A). A set of 5 neuronal types additionally exhibited more than 10 DEGs due to the 5XFAD transgene in either sex. Of these, two cell types (*36_Slc17a6_Nts* and *51_Slc17a6_Ptger3_Gpr83*) exhibited low DEG counts in males at 6 months, but high counts in females of the same age. We thus observed a delayed effect of the transgene on these neurons’ transcriptional profiles in males. In addition, both clusters exhibited high expression of the medial mammillary body markers *Foxb1* and *Rprm* (Supplementary Fig. 9A).

**Figure 6.**
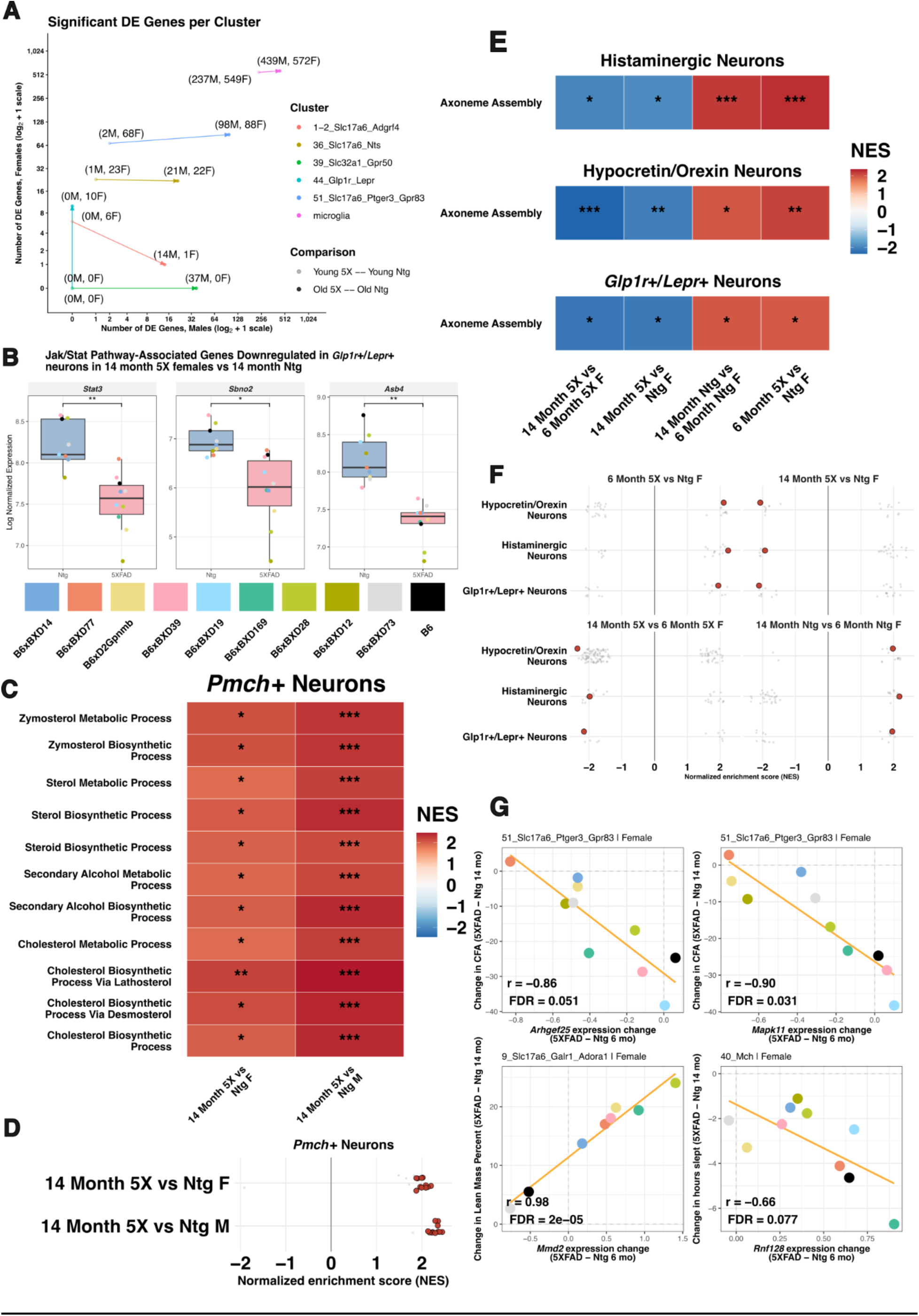
The 5XFAD transgene impacts hypothalamic neuronal types with known roles in memory, energy homeostasis, and circadian rhythmicity. A) Number of DEGs per cluster for males (x-axis) and females (y-axis) between 5XFAD and Ntg conditions at both 6 and 14 month time points. Each line represents a population of cells with at least 10 DEGs in an amyloid vs control comparison; the origin of the line represents the number of DEGs for a comparison at 6 months, and the end indicates the number of DEGs at 14 months. Color of line indicates cluster. B) Stat3, Asb4, and Sbno2 are genes implicated in the Jak2/Stat3 pathway and exhibit lower expression in Glp1r+/Lepr+ neurons from 14 month 5XFAD female mice compared to their Ntg age- and sex-matched counterparts. C and D) Upregulated cholesterol and sterol pathways in Pmch+ neurons between 14 month 5XFAD mice of both sexes compared to their age and sex-matched Ntg counterparts are the pathways with the largest effect sizes. E) Several important neuronal populations involved in sleep and metabolism (Glp1r+/Lepr+ neurons, hypocretin/orexin neurons, and histaminergic neurons) exhibit consistent changes to pathways related to ciliary structure. F) Ciliary pathways denoted in panel E represent some of the most changed (by effect size) pathways in these cell types. G) Relative gene expression changes between 6 month 5XFAD and Ntg female animals of the same strain/age predict differences in 14 month 5XFAD and Ntg phenotypes.

*44_Glp1r_Lepr* (*Glp1r+/Lepr+* neurons) of the arcuate and dorsomedial nuclei and *39_Slc32a1_Gpr50* (GABAergic *Gpr50+* neurons) neurons exhibited divergent sex-specific effects, with *Glp1r+/Lepr+* neurons exhibiting differential gene expression only in females at 14 months and GABAergic *Gpr50+* neurons only in males at 14 months. Neither population exhibited significantly differentially expressed genes at 6 months, indicating the effects of amyloid take longer than 6 months to elicit detectable expression changes at the individual gene level (Fig. 6A). Hypothalamic *Glp1r+/Lepr+* neurons regulate energy balance^43^, metabolism, and satiety, and may mediate in part the brain’s response to semaglutide. *Glp1r+/Lepr+* neurons partly sense nutritional status through activation of the Leptin receptor (*Lepr*) which is mediated by Jak2/Stat3 signaling^44^. Consistent with a dysregulation of Glp1r/Lepr signaling in AD, multiple genes associated with the Jak-Stat pathway (*Stat3*, *Sbno2*, and *Asb4*) were lower in 5XFAD compared to Ntg animals (Fig. 6B).

Changes to genes involved in signaling of *Glp1r+/Lepr+* neurons motivated us to examine whether other physiologically consequential cell types exhibited changes to signaling-related pathways within the hypothalamus. In 14 month 5XFAD vs Ntg comparisons, the strongest detectable signatures in *Pmch+* neurons (involved in coordinating sleep and circadian rhythms) were upregulated cholesterol/sterol synthesis pathways (Fig. 6C, 6D; Supplementary Table 11). Pathways upregulated at 14 months were not significant in 6 month animals (Supplementary Fig. 8B), suggesting an interaction between age and transgene in these neurons.

Many of the strongest pathway results in several physiologically important neuronal types were related to the structure of primary cilia (Fig. 6E,F). Primary cilia are mediators of intercellular signaling within the hypothalamus; some neuropeptide receptors are located on these organelles, and altered function has previously been linked to potent physiological effects^45^. Across three well-studied clusters implicated in either circadian rhythm (*Hcrt+*/hypocretin-orexin neurons and *Hdc*+/histaminergic neurons) or energy balance (*Glp1r+/Lepr+* neurons), there were clear female-specific differences in ciliary assembly between 5XFAD and Ntg animals that occurred in an age-dependent manner: higher expression was observed in 6 month 5XFAD animals when compared to age-matched controls, but at 14 months 5XFAD animals exhibited decreased expression. Furthermore, normal aging in non-transgenic females was associated with increased expression at 14 months compared to 6 month animals; by contrast, in the presence of the 5XFAD transgene, aging was associated with decreased expression (Fig. 6E). Overall, these results indicate that the 5XFAD transgene causes age and sex-dependent changes in expression of ciliary assembly in neuronal clusters that control homeostasis, inverting the age-associated increase observed in female Ntg animals.

We posited that variability in 5XFAD transgene-induced molecular changes may be linked to AD-associated phenotypes, and evaluated whether changes in strain-level phenotypes (defined as the change in mean phenotype between 5XFAD and Ntg mice of the same strain/sex/age from a larger cohort^25^) are associated with altered expression of differentially expressed genes (Supplementary Table 12). In females, we identified several molecular changes at 6 months that predict phenotype change at 14 months, linking early molecular shifts to future phenotypes (Fig. 6G). These include *Mapk11* and *Arhgef25* expression in *Ptger3+/Gpr83+* medial mammillary body neurons linked to associative learning as measured by CFA (greater expression change of *Mapk11* and *Arhgef25* is associated with better cognitive maintenance). The ubiquitin-associated X chromosome gene *Rnf128* in *Pmch+* neurons was associated with sleep time (lower expression change is associated with better ability to maintain sleep time), and *Mmd2* in glutamatergic *Galr1+/Adora1+* neurons was linked to lean mass (lower expression change is associated with better ability to maintain Ntg lean mass levels).

### Conservation of Signatures in the Human HypoAD Dataset

The marker gene *FOXB1* was previously characterized as a conserved, cross-species marker of the medial mammillary bodies^19^. To evaluate whether the changes we observed in these neurons in the 5XFAD model are related to human AD, we performed an integrated clustering analysis of neurons in the hypoAD and ROSMAP MB study, and then grouped clusters with high expression of *FOXB1* together as medial mammillary body neurons. Clusters largely exhibited high expression of either *FOXB1* or *KCNC2*, a lateral mammillary body marker (Supplementary Fig. 9B,C). We performed differential expression analysis separately per dataset to evaluate whether any of the DEGs from the two medial mammillary body neurons in mice (*36_Slc17a6_Nts* and *51_Slc17a6_Ptger3_Gpr83*) from comparisons of 14 month 5X vs 14 month Ntg female mice were associated with AD in human females, after accounting for age and post-mortem interval. Of 85 genes evaluated, 4 genes (*TGFB2*, *RHBDL3*, *MYO16*, and *BMPER*) survived FDR multiple comparisons correction in the HypoAD cohort (Fig. 7 C,D; Supplementary Table 13); no genes were significant in analysis of the ROSMAP MB cohort (Supplementary Table 14), although the direction of association of these genes was consistent.

**Figure 7.**
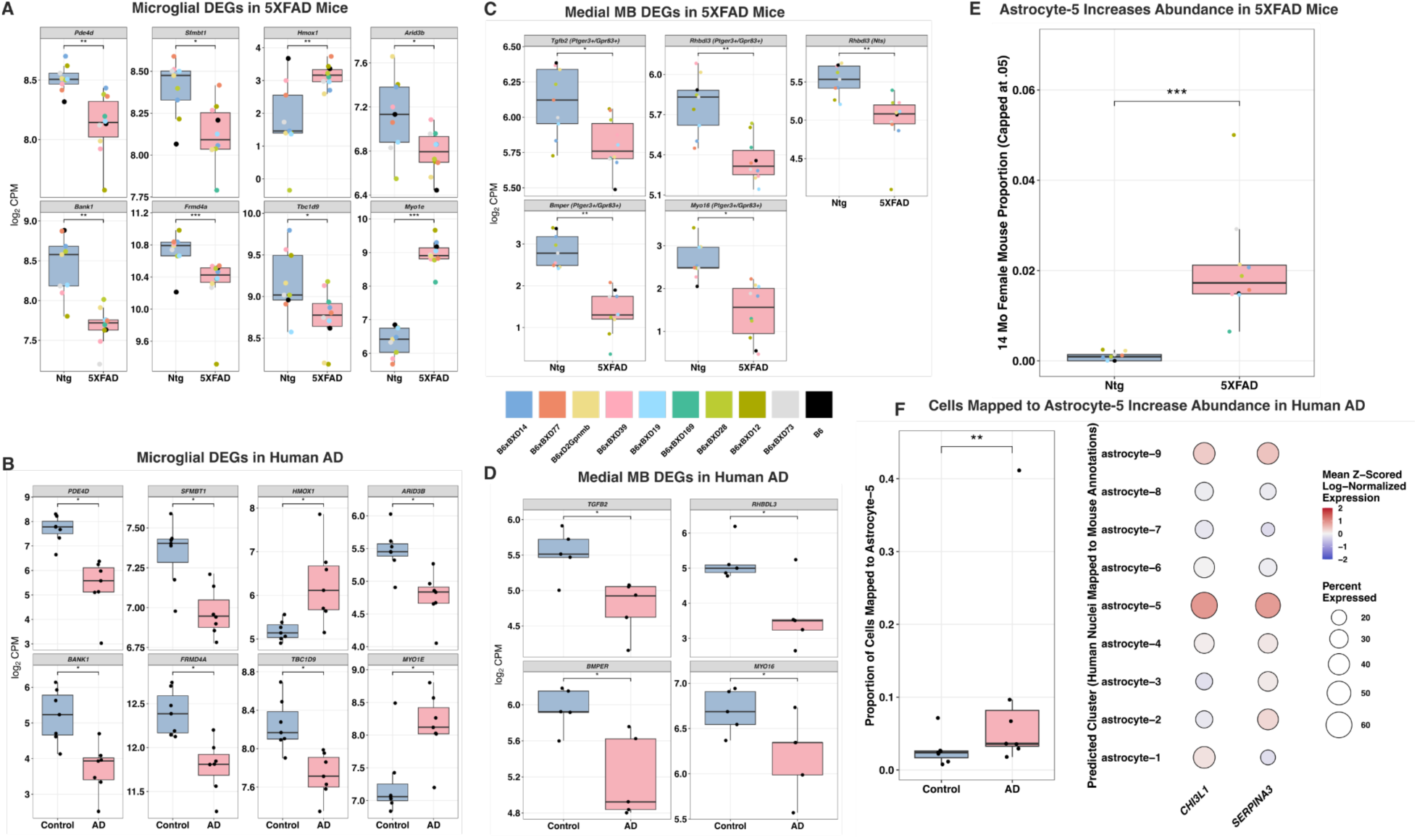
Conservation of amyloid-associated hypothalamic gene expression signatures in the hypothalamus from human AD patients. Conserved microglial (A,B) and medial mammillary body (C,D) DEGs alongside their mouse counterparts. E,F) Human astrocytes that mapped to astrocyte-5 (Serpina3n+/Chi3l1+) via cross-species mapping were more abundant in AD versus non-AD donors, which was consistent with results in mice.

As the hypoAD dataset contains snRNAseq data profiled from three subregions (the mammillary bodies, the arcuate nucleus, and the suprachiasmatic nucleus)^18^, we reasoned this dataset would better reflect identified changes in this study as opposed to the ROSMAP MB study, which profiled a single region. Multiple mouse microglial DEGs retained differential expression signatures in humans using the hypoAD dataset. Of 444 tested associations, a total of 8 genes (*PDE4D*, *SFMBT1*, *HMOX1*, *ARID3B*, *BANK1*, *FRMD4A*, *TBC1D9*, and *MYO1E*) were statistically significant after FDR p-value correction (Fig. 7 A,B; Supplementary Table 15).

We then assessed whether the upregulation of the *Serpina3n+/Chi3l1+* astrocyte-5 cluster in the mouse hypothalamus was conserved in human AD patients via: 1) a cross-species mapping analysis to identify cells that correspond to astrocyte-5 in humans; and 2) differential abundance analysis of these cells. Mapped cells expressed higher levels of *SERPINA3* and *CHI3L1*, orthologs of astrocyte-5 markers in mice, compared to other clusters (Fig. 7F, *right*). We performed the same compositional analysis (consensus of 3 methods, with at least 2 of 3 methods significant) as in mice, and identified a significant association between AD and a higher relative abundance of conserved ‘astrocyte-5’ mapped cells.

## Discussion

This study establishes the hypothalamus as a brain area responsive to dominant genetic mutations that cause early-onset AD. Using high-read depth RNA sequencing, we identified a compendium of cellular and molecular changes across subregions that span the hypothalamus, characterizing areas that have not been adequately examined in previous human AD or animal model studies. To ensure these associations generalize across genetic backgrounds, we profiled 10 strains from the AD-BXD genetic reference panel. This is a notable strength of our study, given that AD therapeutics that are effective in single-strain inbred models often fail to translate to heterogeneous human populations. Incorporating cross-strain associations at the earliest stage of target discovery, as we do here, may improve translatability downstream in the drug discovery pipeline.

We observed several signatures of upregulated neuroinflammation. Both Stage 1 and Stage 2 DAMs and *Serpina3n+/Chi3l1+*-mapped astrocytes were observed at higher levels in transgenic animals, consistent with elevated AD-associated neuroinflammatory signatures in other regions^29,46–49^. In addition, we characterized a sex-biased microglial MHC signature that crossed cluster boundaries. Using the hypoAD dataset, which included human snRNAseq profiles from the suprachiasmatic nucleus, arcuate nucleus, and mammillary bodies, we replicated signatures of neuroinflammation in the 5XFAD model in human AD: elevated relative abundance of *Serpina3n+/Chi3l1+* astrocytes and several microglial DEGs. Conserved microglial DEGs include *HMOX1*, a modulator of inflammation in macrophages^50^ that was previously found at higher levels within microglia around amyloid plaques in other regions in the 5XFAD model^51^, and *FRMD4A.* A prior human genetics study identified a haplotype within *FRMD4A* that was associated with elevated risk of AD diagnosis; the same study characterized several variants of *FRMD4A* associated with the ratio of Aβ42/40, a marker of amyloid progression^52^.

With respect to barrier forming cell types that surround the hypothalamus, several cell types exhibited higher ion transport pathway expression. The concentration of interstitial ions in the brain parenchyma is highly regulated to ensure proper neuronal excitability and glial function, and global concentration change has been identified as a regulator of function^53^. Further, the 5XFAD transgene has previously been shown to decrease K+ clearance of hippocampal astrocytes, which may impact neuronal activity^54^. In addition, a large corpus of work has elucidated relationships between metal ions and the propagation of amyloid and tau^55^; metal ion transport pathways were upregulated in both sexes in endothelial, ependymal, and tanycyte clusters. These findings warrant further investigation into the cellular and molecular consequences of ion dyshomeostasis within the hypothalamus.

Several neuronal cell types linked to cognitive ability, metabolism, and circadian rhythms exhibited molecular changes. With respect to metabolism, these include sex-specific changes to *Glp1r+/Lepr+* neurons in females and *Slc32a1+ Gpr50+* neurons in males. Transcriptional alterations within *Glp1r+/Lepr+* neurons are notable as this cell type is well-characterized for its role in energy homeostasis, and weight loss is a prodromal signature that may precede the onset of cognitive impairment by years. GABAergic *Gpr50+* neurons have previously been localized within the DMH, and Gpr50 was recently deorphanized: the little-LEN (l-LEN) neuropeptide was identified as an endogenous ligand. Activation of Gpr50 impacts metabolism and energy usage, as well as brown adipose tissue-driven thermogenesis^56^. Given the strong influence of these neuronal populations on whole-body metabolism, they may be promising targets to counter the systemic effects on metabolism and weight loss observed in AD. These findings also raise the possibility that the specific cellular and molecular mechanisms mediating metabolic change throughout AD progression are sex-dependent. At the pathway level, we also observed upregulated sterol biosynthesis pathways in *Pmch+* neurons of both sexes at 14 months; *Pmch+* neurons have established roles in energy homeostasis and sleep regulation^57^. Prior studies have linked altered cholesterol with AD progression^58^ including through the AD risk gene *APOE* which is involved in cholesterol transport^59^.

We also characterized unexpected changes in primary cilia assembly expression in cell types that regulate both sleep (orexin/hypocretin and histaminergic neurons) and metabolism (*Glp1r+/Lepr+* neurons). In the hypothalamus, primary cilia are centers of signaling critical to the coordination of energy balance^60^. Multiple neuropeptide GPCRs (including Glp1r^61^, Npy2r^62^, and Mc4r^63^) are located on primary cilia^64^, and several lines of evidence support a role of hypothalamic primary cilia in regulating physiology. Primary cilia length has been explicitly tied to leptin signaling: in a prior study, shortening cilia through siRNA injection of *Kif3a* and *Ift88* within the mediobasal hypothalamus led to a decreased response to leptin alongside other metabolic changes^65^. The length of primary cilia naturally fluctuates with circadian rhythms in neuromedin S-producing neurons; primary cilia in this cell type influence the maintenance of circadian rhythmicity in response to environmental perturbations^66^. Given the influence of primary cilia length on homeostasis, the present work suggests amyloid-induced changes to ciliary structure within the hypothalamus may be a contributor to metabolic or circadian changes observed in AD.

Medial mammillary body neurons are known to be involved in memory formation^67^. In the AD-BXD model, medial MB neurons exhibited the highest DEG count of any neuronal clusters. Four genes (*TGFB2*, *BMPER, RHBDL3*, *MYO16*) exhibited conserved lower expression in humans; notably, *TGFB2* and *BMPER* encode secreted proteins that are members of the TGF-Beta protein superfamily. In an animal model of AD and in cell culture, decreased TGF-Beta signaling in neurons was previously found to increase neurodegeneration and levels of amyloid^68^. The findings here therefore suggest that targeting TGF-Beta in medial mammillary body neurons may be a new avenue to improve memory decline in AD.

In addition to highlighting these findings, we note limitations of the present study. While several associations in comparisons of AD-BXDs versus Ntg controls were conserved in human female AD patients versus controls, the majority of significant DEGs tested in mouse medial MB neurons and microglia did not pass the previously defined significance threshold in humans. One contributor is the limited sample size of available human data used for replication with respect to genetic, environmental, and phenotypic heterogeneity found naturally in human populations; discoveries in larger human studies may lead to the identification of further replicated signatures. With respect to microglia, there are known differences in microglial activation signatures observed through snRNAseq between mice and humans^69^; we thus expected high variability in results. Conserved mammillary body and microglial findings thus represent strong, cross-species signatures of AD within the hypothalamus. Additionally, although we identified several molecular changes that predict future cognitive or homeostatic changes through a strain-level transcriptome-to-phenotype analysis, targeted validation experiments are needed to establish causation between the cellular and molecular changes identified here with non-cognitive phenotypes.

Overall, this study presents a comprehensive resource of the molecular state of the hypothalamus across sex, genetic background, age, and AD transgene status. Our investigation delineated cellular and molecular dysfunction within the hypothalamus, and provides plausible mechanistic explanations for both cognitive (memory) and non-cognitive (sleep, metabolism) symptoms observed in AD. As sleep and metabolic symptoms often precede cognitive impairment and dementia, they represent early causal events contributing to disease pathogenesis; targeting these disease signatures may lead to new therapeutics that are effective either individually or in combination with other AD therapies.

## Acknowledgments

This study was supported by the National Institutes of Aging Grant RF1AG059778, Gene x Environment Interactions in Hypothalamic Dysfunction in Alzheimer’s Disease. Additional support was provided by two grants from the Simons Foundation: SFI-AN-NC-AB-Research-00018041 and Simons Foundation Collaboration on Plasticity in the Aging Brain-984479. J.A. received funding from the Training Grant for Cellular, Molecular and Biomedical Studies (CMBS) at Columbia University Irving Medical Center (5T32GM145766-04). A.R.D received funding from an Alzheimer’s Association Research Fellowship (AARF-18-565506). The authors would like to thank the Single Cell Biology Core at the Jackson Laboratory, which is supported in part by the JAX Cancer Center (P30 CA034196).

The authors acknowledge Doudou Yu and Ashley Webb for providing prepublication access to the hypoAD^18^ dataset.

## Methods

### Discussion of Experimental Design

The 79 animals in our cohort were evenly split across two ages (6 and 14 months), sex (Males and Females), transgene status (maternally derived 5XFAD transgene and nontransgenic controls), and genetic background (10 strains - C57BL/6J (B6), B6xD2Gpnmb, and 8 strains from the AD-BXD panel of genetic diversity). Of note, the DBA/2J genetic background carries a mutation in *Gpnmb*, a gene involved in microglial activation and previously associated with AD; the D2 strain used here (DBA/2J-*Gpnmb*+/SjJ) was edited and matches B6. The 5XFAD transgene is an aggressive model of AD, manifesting its effects through 5 early-onset human AD mutations in *APP* and *PSEN1*. The BXD diversity panel derives from F2 crosses of B6 and DBA/2J (D2) that were then inbred over at least 20 generations.^70^ Therefore, all BXD animals are related, but individual strains retain distinct genotypes. The AD-BXD panel consists of an F1 cross between a male mouse of one of the BXD strains, crossed with a female B6-5XFAD mouse (a B6 mouse with the 5XFAD transgene).^24^ Consequently, each animal contains one allele of each chromosome that is largely the same, from the B6-5XFAD, and one that is genetically distinct, from the nontransgenic BXD.

### Mouse Line Generation, Husbandry, and Phenotyping

AD-BXD mice were generated as described previously by crossing female B6-congenic mice hemizygous for the 5XFAD transgene with males from 10 BXD strains^24,71^. Mice were maintained on a 12 h/12 h light/dark cycle. Mice were group-housed and fed ad libitum a normal control diet (LabDiet Diets 5K0G). All animal experiments were performed in accordance with the NIH Guide for the Care and Use of Animals and were approved by the Institutional Animal Care and Use Committee at The Jackson Laboratory.

Mice underwent a series of cognitive and sensorimotor phenotyping. Body weights were measured every two months, starting at 2 months of age. Mice were tested using the y-maze assay every 2 or 4 months throughout their lives, starting at 2 months of age. At 3 months, 6 months, or 14 months, mice underwent sensorimotor testing (grip strength, inclined screen, and narrow beam tests) and contextual fear conditioning (CFC). A measure of short term memory acquisition, contextual fear acquisition (CFA), was assessed as the percent freezing in the post shock four period of CFC. Long-term memory was assessed via contextual fear memory (CFM). Mice were habituated to transport and testing for two days prior to behavioral testing. Additionally, a subset of mice of the same strains (not assessed with snRNAseq in this study) were longitudinally tested with nuclear magnetic resonance body composition analysis, glucose tolerance testing, and indirect calorimetry. Details on cognitive and sensorimotor phenotyping are described elsewhere.^25^

### Tissue Collection

Immediately following CFC, mice were euthanized using isoflurane and brains were removed. Hypothalami were dissected and flash-frozen in liquid nitrogen for subsequent biochemical analysis.

### Nuclei isolation and sequencing

Nuclei were isolated from snap frozen brain fragments according to an in-house developed protocol leveraging the gentleMACS tissue dissociator system (Miltenyi Biotec)^72^. Briefly, tissue fragments were dissociated in ice cold Nuclei Isolation Buffer comprising nuclei extraction buffer (Miltenyi Biotec, 130-128-024) and RNAse inhibitor (Millipore Sigma, 3335399001) using the 5-minute ‘4C_nuclei’ protocol provided by Miltenyi. The protocol was customized to add additional mixing time based on tissue fragment mass. Resulting digests were filtered through a 100um filter and pelleted in cold PBS + 2% BSA + 1mM DTT + 1U/ul RNAse inhibitor. Depending on the level of debris, nuclei suspensions were further purified by additional washes or via sucrose column purification.

Nuclei concentration and viability was assessed on a LUNA FX7 automated cell counter (Logos Biosystems), and up to 40,000 nuclei from each suspension were loaded onto one lane of a 10x Genomic Chip M. Single cell capture, barcoding and library preparation were performed using the 10x Genomics Chromium X platform version 3.1 NEXTGEM HT chemistry and according to the manufacturer’s protocol (#CG000416)^73^. cDNA and libraries were checked for quality by Tapestation 4200 (Agilent) and Qubit Fluorometer (ThermoFisher), quantified by KAPA qPCR, and sequenced at 25% (1 lane) of an Illumina NovaSeq 6000 S4 v1.5 200 cycle flow cell lane, with a 28-10-10-90 asymmetric read configuration, targeting 20,000 barcoded nuclei with an average sequencing depth of 120,000 reads per nucleus.

### Read Alignment

Illumina base call files for all libraries were converted to FASTQs using bcl2fastq v2.20.0.422 (Illumina) and FASTQ files associated with the gene expression libraries were aligned to a customized reference assembly containing GRCm39 and 5XFAD assemblies and annotations using CellRanger count version 7.0.0 (10x Genomics).

### Quality Control (QC) and Initial Clustering

Decontamination analysis was performed for each library separately using Decontx^74^ with default parameters except for the z parameter, which takes in cell labels to assist in decontamination. Preliminary broad cell type classifications were used as the cell labels, which were determined via Seurat’s sketch clustering workflow^75^ followed by marker gene analysis. Doublet detection was then performed on each library separately using scDblFinder^76^ with default parameters on all cells with greater than 500 UMIs. Cells were removed if they were identified as doublets.

Cells were filtered out if mitochondrial expression was greater than five percent; if the number of unique genes per cell was outside of the range 500 - 10,000; or, if the number of UMIs was outside the range of 2,000 - 50,000. Taking inspiration from a recently published technique to assess QC for scRNAseq based on genes correlated with *Malat1* expression (a predominantly cell nucleus-localized marker gene), we calculated a QC score for snRNAseq composed of summed log1p normalized expression of 61 genes^26^. We filtered out nuclei if their QC score values were greater than 100 (this threshold was chosen after examining the distribution of scores).

Gene expression profiles were then initially clustered using Seurat’s sketch clustering workflow^75^. First, log-normalization and variable feature identification was performed on the entire dataset. A subset of 100,000 cells was selected using the leverage score method. On this subset, we reidentified variable features, scaled log-normalized expression, ran principal components analysis (PCA), found nearest neighbors using the top 50 dimensions, and performed clustering using the Louvain algorithm with a resolution of 2×10^−3^. We assigned names to these broad cell classifications by identifying marker genes via the FindAllMarkers function. Cell class assignments were then projected to the rest of the dataset using the ProjectData function.

Counts were changed from ENSEMBL IDs to gene symbols using the AnnotationDBI^77^ and org.Mm.eg.db^78^ packages on Bioconductor; the mapIDs function was used to convert between the two nomenclatures using the following parameters: column = “SYMBOL”, keytype = “ENSEMBL”, and multiVals = “first”. ENSEMBL IDs that did not map to a gene symbol were removed. If multiple ENSEMBL IDs corresponded to the same gene symbol, then the UMIs for the non-ambiguous gene symbol for each cell were determined by taking the max value of all the ENSEMBL IDs of that cell.

### Neuronal and Glial Subclustering

Neuronal gene expression values were log-normalized and the top 3000 variable features were identified using Seurat’s FindVariableFeatures function. Gene expression was subset to just these variable features, which were scaled, and the top 50 principal components (PCs) were identified using BPCells. Using the PCs, we produced a graph containing the top 20 k-nearest neighbors (KNN) for each cell; this graph was then converted to a shared nearest neighbors (SNN) graph. Leiden clustering was performed on the SNN graph using the cluster_graph_leiden function from the BPCells^79^ package (which itself employs the cluster_leiden function from the igraph^80^ package), with the objective function parameter set to the default, “modularity”, and the resolution set to 0.5. Small clusters with less than 100 cells were removed. Marker genes were identified using log-normalized expression of each cluster using the marker_features function from BPCells. Cluster quality was evaluated with a number of metrics, including: pairwise modularity ratio, neighborhood cluster purity, silhouette score (calculated on a 5% subsample of cells for all clusters with more than 1000 cells), and a pairwise correlation heatmap displaying the Spearman correlation between clusters.

All neuronal clusters with more than 20,000 cells were selected for subclustering using largely the same steps as above except the number of variable genes identified was 2000; the number of PCs identified was 30; and the Leiden clustering resolution was set to 0.1; however, if the modularity score of the subclustering was not greater than 0.95, the cluster was left as is. Clusters with fewer than 100 cells were removed. Silhouette scores and cluster quality metrics were calculated across all clusters, and any cluster with a median silhouette score below 0 was merged with its closest cluster (closest cluster determined by Spearman correlation of PCs). Cluster quality metrics were then determined again. If any clusters that were not subclustered previously exhibited a median silhouette score less than 0, it was merged with its closest cluster (based on Spearman correlation of PCs), if the correlation with its closest cluster was greater than 0.5.

Magnocellular *Avp+/Oxt+* neurosecretory cells were segregated in the initial Seurat clustering, and were subclustered separately than other neuronal clusters. A Leiden resolution of 0.05 was used. Clusters with less than 500 cells were removed and clusters with a median silhouette score (as determined by a subsample of 10% of each cluster) below 0 were merged with its most similar cluster. Marker gene expression was used to distinguish *Avp+ vs Oxt+* magnocellular neurons.

Glial class groupings from the initial class assignments (astrocytes/ependymal/tanycytes, microglia/immune/border associated macrophages (BAMs), oligodendrocytes, oligodendrocyte precursor cells (OPCs), vascular, and vascular and leptomeningeal cells (VLMCs)) were subclustered separately. A cluster corresponding to the pituitary classes (ex: somatotrophs) was removed, as these cells likely reflect contamination from the pituitary and are not present in the hypothalamus.

Astrocyte/ependymal/tanycyte cells were initially clustered together. PCs were found using the top 2000 variable genes as above, followed by nearest neighbor determination and leiden clustering on the shared nearest neighbor graph with a resolution parameter of 0.1. Clusters with less than 500 cells were removed, and doublet clusters were manually identified and removed based on marker gene expression. The first round of clustering distinguished astrocytes, ependymal, and tanycyte cell classes. Each of these cell classes was further clustered individually following the same parameters, and clusters with a median silhouette score (as determined by a subsample of 10% of each cluster) below 0 were merged with its most similar cluster. One tanycyte cluster was manually removed after this step because it expressed high levels of ependymal markers. Most astrocytes belonged to a single cluster, which was subsequently subclustered.

Microglia, BAMs, and immune cells were initially grouped into one class. PCs were found using the top 2000 variable genes as above, followed by nearest neighbor determination and leiden clustering on the shared nearest neighbor graph with a resolution parameter of 0.1. Clusters with less than 500 cells were removed, and doublet clusters were manually checked, identified, and removed based on marker gene expression. T-cells, microglia, and BAMs were distinguished. Microglia were subclustered further following the same steps, and clusters with a median silhouette score (as determined by a subsample of 10% of each cluster) below 0 were merged with its most similar cluster. Two subsequent rounds of iterative clustering were performed for the microglia. For the microglial subclustering rounds, multiple Leiden resolutions were tested and one was manually chosen based on modularity and effectiveness of the clustering to distinguish a substantial fraction of cells.

Clustering for the oligodendrocyte, OPC, vascular, and VLMC cell classes was performed in the same manner. PCs were found using the top 2000 variable genes as above, followed by nearest neighbor determination and leiden clustering on the shared nearest neighbor graph with a resolution parameter of 0.1. Clusters with less than 500 cells were removed, and doublet clusters were manually checked, identified and removed based on marker gene expression. After doublet removal, the remaining cells in each class were subclustered as above. Clusters with a median silhouette score (as determined by a subsample of 10% of each cluster) below 0 were merged with the most similar cluster.

### Neuronal Cluster Nomenclature (Identification of Unique Neuropeptide/Receptor Marker Genes or Map to Established Types)

We first stratified clusters by their expression of the canonical neurotransmitter transporter genes *Slc17a6* (excitatory transporter) and *Slc32a1* (inhibitory transporter) by setting a cutoff of 25% expression in a cluster. All clusters expressed either *Slc17a6* or *Slc32a1* except for one cluster that expressed both.

Building off a prior gene list^81^, we produced a gene set containing neuropeptide/neuropeptide-receptor genes, and used an additional list of GPCRs (Gene Ontology/GO term 0004930, which is described as a G protein-coupled receptor activity gene set^82^. We filtered out from the GO set any terms that contain the following words in their description - “vomeronasal”, “taste”, “predicted”, and “olfactory”. We included the additional set of GPCRs to provide a more comprehensive identification of signaling-related marker genes; for example, orphan GPCRs that do not yet have known, characterized agonists.

Neuronal nuclei were counts per million (CPM)-normalized across all genes, and then subset to the aforementioned genes. We then performed mean-pseudobulk aggregation to get the average CPM expression per cluster across the dataset. We binarized expression, so each cluster was noted as either expressing a gene or not, according to these rules: 1) 25% of the max gene-expressing cluster is set as the binary expression threshold unless 2) this value is greater than 100 CPM, in which case the threshold is set to 100 CPM, or 3) this value is less than 10 CPM, in which case the threshold is set to 10 CPM.

We next took inspiration from an algorithm presented in a BICCN whole brain transcriptome profiling study to assign names to neuronal clusters, where marker genes were identified via mixed integer linear programming (MILP) although not at the level of specificity we do here on a pre-defined biological gene set^81^. We used MILP to identify the minimal set of neuropeptide, receptor, or GPCR genes that distinguish each cluster from every other cluster of the same neurotransmitter type. We implemented this algorithm in R via the ompr package^83^, and used the glpk (GNU Linear Programming Kit) solver^84^. For the one cell type expressing both neurotransmitters, we identified a unique gene set combination across all clusters.

If a cluster expressed a previously well-characterized combination of neuropeptides or receptors, we assigned those genes as its name to be consistent with prior literature. For these established clusters, the nomenclature scheme was to add an internal number we used to identify clusters before assignment, followed by the established name (ex: 44_Glp1r_Lepr). For the majority of the clusters which did not have well-established names, our nomenclature scheme was to first identify the cluster by an internal number we used to identify clusters before assignment, followed by the cluster’s neurotransmitter and the unique neuropeptide/receptor combinations (ex: 35_Slc17a6_Gpr12_Gpr171). For cluster 51, one of the medial MB neuronal clusters mentioned in Fig. 6, we manually assigned the unique neuropeptide/receptor combination to *Ptger3+/Gpr83+*.

### Identification of Genes Influenced by Genetic Background

We identified genes influenced by genetic background for each cell class separately; neurons were grouped as either GABAergic or glutamatergic based on whether they expressed *Slc17a6* or *Slc32a1* past the 0.25 fraction expressing threshold. We removed one cluster that expressed both transporters from this analysis. We first removed all gene models and RIKEN genes from the gene expression matrix. For each cell class separately, we: 1) determined the top 5000 variable genes and subsequently subset to these features; 2) performed a summed pseudobulk, or aggregated expression over all cells by adding expression 3) performed log CPM normalization; 4) assessed the relative contribution of each of the variables in this dataset (genetic background, sex, age, transgene), treating each as a random effect; 5) used the fitExtractVarPartModel^85^ function from the VariancePartition package to estimate contributed variance of each cluster for each variable gene. The Nelder-Mead optimizer^86^ was used for fitExtractVarPartModel. We note that the method of variable gene selection, which uses a variance-stabilizing transform (VST) after log-normalization, is non-standard as a VST is typically applied to raw counts. A recent benchmark identified that variable gene selection via VST on log-normalized data outperforms VST on raw counts.^87^

### Microglial Factor Decomposition and Downstream Analyses

Microglia from animals with less than 1000 cells profiled or without CFA/CFM profiling were excluded from this analysis. Briefly, the top 3000 variable genes were identified, and microglial gene expression was subset to these genes. De novo factor discovery was performed using sciRED^32^, which accounts for differences in read depth between cells via Poisson regression, performs PCA on resulting pearson residuals, and rotates PCs to produce more interpretable factors.

Factor scores per animal (for all animals with greater than 50 cells profiled, otherwise excluded) were calculated by averaging across all cells, before linear modeling using the formula “∼ age_genotype + age_genotype:sex + 0 + (1|strain)” and statistical testing of specific contrasts/comparisons using the glht function from the multcomp package^88^. Multiple comparisons FDR correction was performed for each contrast separately, across the 50 factors tested for that contrast.

### Visualization of Microglia Expression via UMAP

UMAP^89^ was used to visualize microglial gene expression in two dimensions. Briefly, the top 30 principal components were identified based on the scaled, log-normalized expression values of the top 2000 variable genes using BPCells^79^. UMAP coordinates were calculated from PCs using the uwot function (from the umap^90^ package in R).

### Differential Gene Expression Analysis

Differential expression analysis was performed using dreamlet^91^, which enabled us to perform mixed modeling on pseudobulked data. The single animal without CFA or CFM measurements was excluded from this analysis. Analysis was performed with all neuronal cell types at the cluster level, and glia at the cell class level; given their unique function and attributes^34^, GLS astrocytes were grouped separately than other astrocytes for this analysis. Gene models and RIKEN genes were removed prior to differential analysis by filtering out genes that start with “Gm” or “Rik”.

Multiple contrasts were tested for each sex separately, and named according to the comparison being performed: 1) 14 month 5X vs 14 month Ntg (prolonged_amyloid_contrast_m or prolonged_amyloid_contrast_f); 2) 6 month 5X vs 6 month Ntg (early_amyloid_contrast_m or early_amyloid_contrast_f); 3) 14 month Ntg vs 6 month Ntg (aging_contrast_m or aging_contrast_f); 4) 14 month 5X vs 6 month 5X (late_vs_early_amyloid_contrast_m or late_vs_early_amyloid_contrast_f). Contrasts between sexes were also performed: 1) 6 month Ntg female vs 6 month Ntg male (effect_of_sex_young_ntg); 2) 6 month 5XFAD female vs 6 month 5XFAD male (effect_of_sex_young_5x); 3) 14 month Ntg female vs 14 month Ntg male (effect_of_sex_old_ntg); 4) 14 month 5XFAD female vs 14 month 5XFAD male (effect_of_sex_old_5x).

Animals were annotated by their ID, age, transgene status, strain, and sex. The formula “∼ age_genotype + age_genotype:sex + 0 + (1|strain)” was used in the processed_assays and dreamlet functions, which accounts for strain as a random intercept. We used a workflow consisting of the functions aggregateToPseudobulk, processAssays, and dreamlet. Multiple comparisons correction was performed across genes for each cluster and contrast separately using the FDR method.

### Consensus Differential Relative Abundance Analysis

Differential composition analysis evaluating changes to the relative abundance or proportion of a given cluster, relative to other clusters within the same cell class, was performed through a consensus approach that combined the results of three methods: sccomp^30^, ANCOM-BC2^92^, and voomCLR^31^. We reasoned that using a consensus approach, requiring that at least 2 of 3 methods converged on a biological finding (FDR-corrected p-value less than 0.10, and same direction of association) to call an association, would improve robustness. This is important as methods use different underlying algorithms and can produce variable results when applied to the same sc/snRNAseq datasets^30^; therefore, relying on a single differential composition method may lead to conclusions that are not robust.

Datasets were preprocessed in the same way for all methods: animals with fewer than 1000 cells were excluded, as was the single animal without CFA or CFM behavioral measurements. Cell classes with only one cluster (ex: T-cells or BAMs) were removed.

For all methods, we accounted for strain as individuals with the same genetic background are not independent observations. Due to method specification differences (ex: voomCLR does not natively allow a random intercept term for strain, and ANCOM-BC2 does not allow a term that denotes an intercept of zero), we accounted for non-independence through somewhat different modeling approaches and statistical contrasts for each method. For all methods, statistical tests were performed to evaluate the effects of the 5XFAD transgene on cluster relative abundance at 14 months and 6 months compared to their same age/sex Ntg counterparts; the effects of mid-life aging on cluster relative abundance (comparing 14 month to 6 month per genotype/sex); and, the effects of sex on cluster relative abundance (comparing males to females per age/genotype).

*Sccomp* analysis: The sccomp^30^ workflow consists of running the sccomp_estimate (to model relative abundance), sccomp_remove_outliers (to remove outliers), and sccomp_test (to perform hypothesis testing) functions. Animals were given a designation by their age and genotype (age_genotype). The mixed effects formula “∼ age_genotype + age_genotype:sex + 0 + (1|strain)” was used to model composition; strain was used as a random intercept to account for non-independence of observations from the same genetic background. Count relative abundance was modeled through the sccomp_estimate function, and default parameters were used except the bimodal_mean_variability_association parameter was set to True. sccomp_remove_outliers was used with the percent_false_positive parameter set to 10. Contrasts were evaluated using the sccomp_test function with the test_composition_above_CLR_fold_change parameter set to 0. Adjusted p-values provided directly by sccomp (in the c_FDR column) were used to assess statistical significance.

*ANCOM-BC2*^92^ analysis: Animals were given a designation of age/genotype/sex prior to modeling. The ancombc2 function was used to model composition, with the age/genotype/sex designation per animal set as the fix_formula parameter and strain set as a random intercept through the rand_formula parameter. Multiple comparisons FDR correction was performed across clusters for each class/contrast separately.

*VoomCLR*^31^ analysis: Animals were designated by their age and genotype, and the voomCLR function was run on counts using a design matrix created via the formula “∼ age_genotype + age_genotype:sex + 0” with default parameters, except the varCalc parameter was set to “analytical”. We followed instructions from the package vignette discussing how to account for non-independence. Specifically, we: 1) used the duplicateCorrelation function from limma; 2) reran the voomCLR function given these results; and 3) ran duplicateCorrelation a second time given the resulting voomCLR output. The lmFit and eBayes functions were then used to fit a linear model and produce moderated statistics via empirical Bayes shrinkage; p-values and effect sizes for each statistical contrast were then acquired sequentially via the topTableBC function with the bootstrap parameter set to “parametric”, the contrastMatrix parameter subset for each contrast; and, the voomWeights parameter set using the output of voomCLR. Multiple comparisons FDR correction was then performed across clusters, separately for each cell class/statistical contrast.

### Identification of Clusters that Exhibit Compositional Associations with Strain (Relative to C57BL/6J)

We identified clusters with compositional changes, relative to its cell class, that are different from B6 using the consensus differential abundance method largely as described previously. The formula “∼ strain + age + sex + genotype” was used for all three methods. Given each variable was treated as a fixed effect, the duplicateCorrelation function and other steps to account for non-independence were not run for voomCLR; additionally, the topTableBC function in voomCLR was used with default parameters.

### Transcriptome-to-Phenotype Strain-Level Analysis

Gene Expression: Log1p-normalized expression per cluster was calculated for significant differentially expressed genes between 5XFAD and Ntg animals of the same strain/sex/age. Per-strain differences between 5XFAD and Ntg animals for each gene were determined by subtracting the log1p expression of the Ntg animal from the 5XFAD animal.

Phenotyping: Phenotyping metrics were determined at the strain level by averaging across mice of a larger cohort of AD-BXDs of the same age/sex/genotype/strains^25^ used in this study, only including mice fed a chow diet to match the diet of the animals profiled with snRNAseq. Change in phenotype was determined using Ntg animals as the reference group. Raw differences in CFA, sleep time, lean mass percentage, fat mass percentage, and a z-scored sensorimotor composite score were used as phenotypes; percent weight change was calculated using the Ntg group as the reference.

We filtered out animal/cluster pairs where there were less than 30 cells profiled, and required at least 8 strains (out of a maximum of 10) with both Ntg and 5XFAD animals to meet this threshold. Pearson correlation was used to evaluate the relationship between strain-level expression change and strain-level phenotypes. We tested the combinations of 6/14 month expression change (using DEGs specific to a time point) and 14 month phenotype change between AD-BXDs and Ntg animals. FDR multiple comparisons correction was performed across significant DEGs for each cluster/sex/phenotype/combination of expression change and phenotype change timepoints separately, using a predefined threshold of 0.10 for significance given the limited number of strains profiled.

### Gene Set Enrichment Analysis (GSEA)

Gene set enrichment analysis was performed on dreamlet results using the gseGO function from the clusterProfiler^93^ package. We reran dreamlet at the cluster level for barrier cell types (vascular, VLMC, ependymal, and tanycyte clusters) prior to GSEA, since the initial dreamlet analysis was performed considering all cells of a class together.

The following parameters were used in the gseGO function: OrgDb = “org.Mm.eg.db”, ont = “BP”, keyType = “SYMBOL”, pAdjustMethod = “BH”, pvalueCutoff = 1, by= “fgsea”, nPermSimple = 50000, seed = TRUE, and eps = 1 x 10^−50^. FDR-provided p-values were used to assess statistical significance.

### Selection of Ion Transport Pathways in Figure 5 and Supplementary Figure 7

Ion transport pathways shown in Figure 5 and Supplementary Fig. 7 are either GO terms 0006811 (monoatomic ion transport)/0015698 (inorganic anion transport) or a descendant of these terms that exhibited statistically significant changes in comparisons of 14 month 5XFAD vs Ntg of either sex for a given cluster. Descendant terms with “regulation” or “Regulation” were excluded. Highlighted comparisons in Figure 5 heatmaps are a subset of these clusters/comparisons.

### Integrated Rosmap MB and HypoAD Medial MB Cluster Assignment and Differential Expression Analysis

Rosmap MB libraries were aligned using starSolo^94^ with the following parameters: runMode = “alignReads”, soloType = “CB_UMI_Simple”, soloUMIlen = 12, soloFeatures = GeneFull_Ex50pAS, soloCellFilter = EmptyDrops_CR, soloCBmatchWLtype = 1MM_multi_Nbase_pseudocounts, soloUMIfiltering = MultiGeneUMI_CR, soloUMIdedup = 1MM_CR, clipAdapterType = CellRanger4, and outFilterScoreMin = 30. Read counts were fed into Decontx^74^ decontamination analysis followed by scDblFinder^76^ for doublet removal. HypoAD cell counts from the mammillary bodies and Rosmap MB counts were then filtered for the following QC metrics: total counts between 2000 and 100,000; mitochondrial read percent less than 5 percent; non-doublets; and, for hypoAD, primary data. Rosmap MB nuclei were additionally filtered to have at least 500 genes detected.

Principal components analysis was performed to identify the top 30 PCs of the combined matrix using the top 2000 variable genes. Harmony batch correction^95^ was then used to correct for differences between datasets. Broad clusters were acquired via creation of a shared nearest neighbors graph, then Leiden clustering using a resolution of 0.01. Non-doublet neuronal clusters were grouped together into one class.

Just on the neuronal counts matrix, principal components analysis followed by Harmony integration across datasets were then performed, followed by clustering on the shared, batch-corrected embedding with a Leiden resolution of 0.1. Neuronal clusters 3 and 5 were assigned as medial MB neuron clusters based on marker gene expression, and then dreamlet differential expression analysis was performed per-dataset to identify the main effect of AD diagnosis after accounting for pmi and age (pmi and age values were scaled relative to the values of other donors in a given dataset). Results were subset to the human orthologs of genes that were significant in either of the two medial MB clusters in the mouse dataset in the 14 month 5X vs 14 month Ntg female comparison. Two-tailed p-values were converted to one-tailed p values based on the direction of mouse association, followed by FDR correction. If any gene was significant in both mouse clusters but in a different direction of association, it was excluded.

### HypoAD Microglia DEG Analysis

The hypoAD dataset was subset to microglia, followed by PCA (performed on top 2000 variable genes) and batch correction using harmony in order to correct across hypoAD batches. Leiden clustering, performed on a shared nearest neighbors graph and using a resolution of 0.1, was used to identify cell subgroups. One cluster expressed marker genes indicative of border-associated macrophage marker genes; this cluster was removed. Differential expression analysis was then performed on the remaining microglial cells using dreamlet as described for the medial MB neurons.

### HypoAD Astrocyte Mapping and Differential Abundance Analysis

Astrocytes from the hypoAD dataset were combined in a matrix with astrocytes from the mouse AD-BXD dataset; both datasets were subset to genes with 1:1 mouse to human orthologs. PCA was performed on the top 5000 variable genes, followed by Harmony correction across species to create an integrated embedding.

Multiple machine learning models were next evaluated for their accuracy to assign mouse cells to mouse clusters on this embedding, before applying these models to assign human cells to mouse clusters. First, the mouse cells were split into an 80%/20% train/test set, stratified by cluster; a set of 5000 cells per cluster were sampled from the train set for model training. If less than 5000 cells were present for a given cluster, synthetic data for that cluster was generated using the synthetic minority over-sampling technique (SMOTE)^96^. XGBoost^97^, random forest, and ridge regression models were trained using this training set and model evaluation was performed using the test set. The F1 score was the highest for XGBoost. An XGBoost model was then trained using a sample of the entire mouse astrocyte dataset (5000 cells per cluster; SMOTE used to generate synthetic data if less), followed by cluster assignment. Differential relative abundance analysis was performed using the consensus of 3 methods largely as previously described on the mapped clusters, accounting for age of death and postmortem interval as covariates (formula for all three methods: “∼ ad + age + pmi_hours”). As this was a prespecified targeted analysis, p-values from voomCLR and ANCOM-BC2 were not adjusted; the FDR value returned by sccomp was used to determine consensus.

### Validation of Upregulated MERFISH Stage 1/2 DAM Levels

An anndata object containing the MERFISH imaging spatial transcriptomics dataset was acquired from the Brain Image Library.^98^ The MERFISH dataset was subset to hypothalamic microglia using previously assigned region and cell type assignments. Genes were filtered to only include those with more than 50 cells with detectable expression, and cells were filtered to include only those with at least 100 transcripts of this gene subset. The snRNAseq dataset in the present study was filtered to only microglia, and to shared genes. The MERFISH and snRNAseq datasets were concatenated, and the top 20 PCs were identified. Harmony was then applied to PCs to account for differences between the MERFISH and snRNAseq methods.

Three models (random forest, glmnet, and xgboost) were then assessed for performance to classify snRNAseq microglia to the appropriate cluster using these embeddings, prior to classification of the MERFISH cells. The microglia snRNAseq matrix was split into an 80/20 train/test set, and each cluster in the training set was subset to 5000 cells; if less than 5000 microglia were present, SMOTE was used to create synthetic observations of that cluster to reach 5000 cells, prior to model training and evaluation on the test set. Random forest was chosen based on F1 score. A random forest model was then trained on a dataset consisting of 5000 cells per cluster from the snRNAseq embeddings (upsampled with SMOTE if less than 5000 cells) and used to classify MERFISH cells. Consensus differential abundance was performed largely as described previously to evaluate associations with the 5XFAD transgene. Hemizygous 5XFAD and 5XFAD animals were grouped together based on 5XFAD status, and statistical models accounted for sex as a main effect (formula for all three methods: “∼ 5XFAD status + Sex”). Sccomp FDR p-values were acquired directly from the method, and voomCLR/ancombc2 p-values were converted to one-tailed p-values to assess the previously identified increase in relative abundance, and adjusted using Bonferroni correction.

## Generative AI Disclosure

Large language models were used for code and debugging assistance.

## Data and Code Availability

Raw sequencing data is available on SRA (accession: PRJNA1251037). Processed snRNAseq data from this study is available on the CZI CellxGene Explorer^99^ at https://cellxgene.cziscience.com/collections/7039e317-12d2-424a-8ea6-479d6e7a8cf0. Code to reproduce figures/analyses are located on Github at https://github.com/jonathan-algoo/hypothalamus-adbxd-atlas. An additional repository containing code, metadata, datasets used in analysis (besides the ROSMAP mammillary body dataset which is not distributed due to data sharing restrictions), and instructions to reproduce analyses/visualizations is stored on Hugging Face at: https://huggingface.co/datasets/jonathan-algoo/hypothalamus-adbxd-atlas.

For access to the ROSMAP mammillary body dataset, due to data control regulations, we refer readers to the original publication.^19^

## Supplementary Table Legend

**Supp. Table 1** Genes, per broad cell type, where 30% or more of total variance was attributed to genetic background.

**Supp. Table 2** Number of variable genes due to genetic background per broad cell type.

**Supp. Table 3** Results from voomCLR, sccomp, and ANCOM-BC2 differential relative abundance analyses.

**Supp. Table 4** Microglial Factor Decomposition - Gene x Factor Loadings

**Supp. Table 5** Microglial Factor Decomposition - Cell x Factor Scores

**Supp. Table 6** Significant amyloid related associations from microglial factor analysis.

**Supp. Table 7** Significant dreamlet differential expression analysis results (neurons at cluster level, glia at broad cell type level).

**Supp. Table 8** GSEA pathway analysis results for GLS astrocytes.

**Supp. Table 9** Significant barrier celltype cluster-level differential expression analysis results.

**Supp. Table 10** Significant barrier celltype cluster-level GSEA pathway analysis results.

**Supp. Table 11** Significant results from GSEA pathway analysis of well-known/selected neuronal clusters.

**Supp. Table 12** Significant results from transcriptome-phenotype analysis at the strain level.

**Supp. Table 13** Replication analysis using dreamlet - medial MB DEGs in humans (hypoAD).

**Supp. Table 14** Replication analysis using dreamlet - medial MB DEGs in humans (Rosmap MB).

**Supp. Table 15** Replication analysis using dreamlet - microglial DEGs in humans (hypoAD).

## Supplementary Figures

**Supplementary Figure 1.**
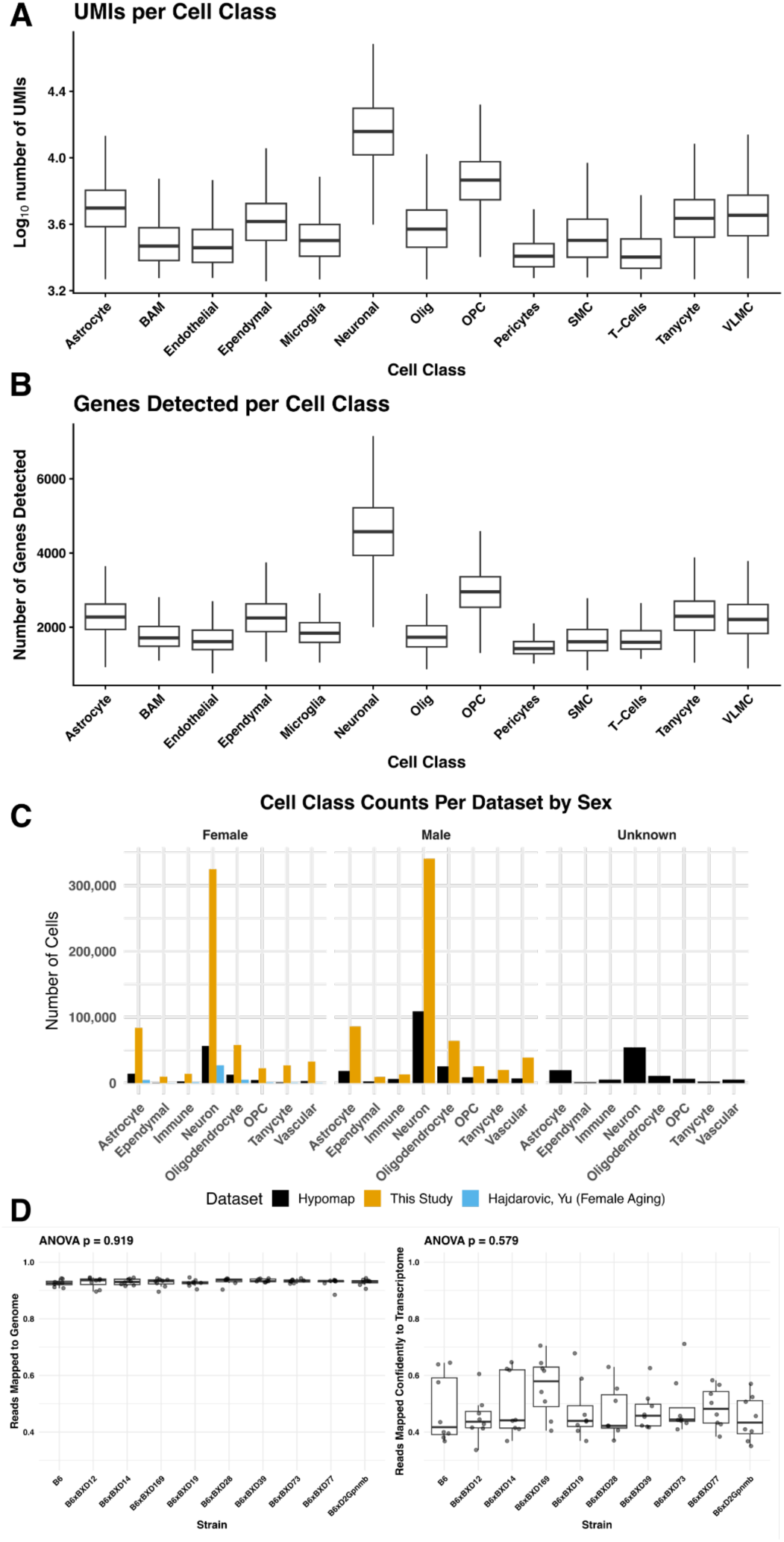
Descriptive statistics and QC metrics: a) Number of UMIs (log_10_ scale), stratified by cell class. b) Number of genes detected per cell class. c) Number of cells per cell type relative to other datasets.^100,101^ d) Cellranger quantification of reads mapped to genome and e) reads mapped confidently to transcriptome (right) indicate similarities in alignment performance across strains.

**Supplementary Figure 2.**
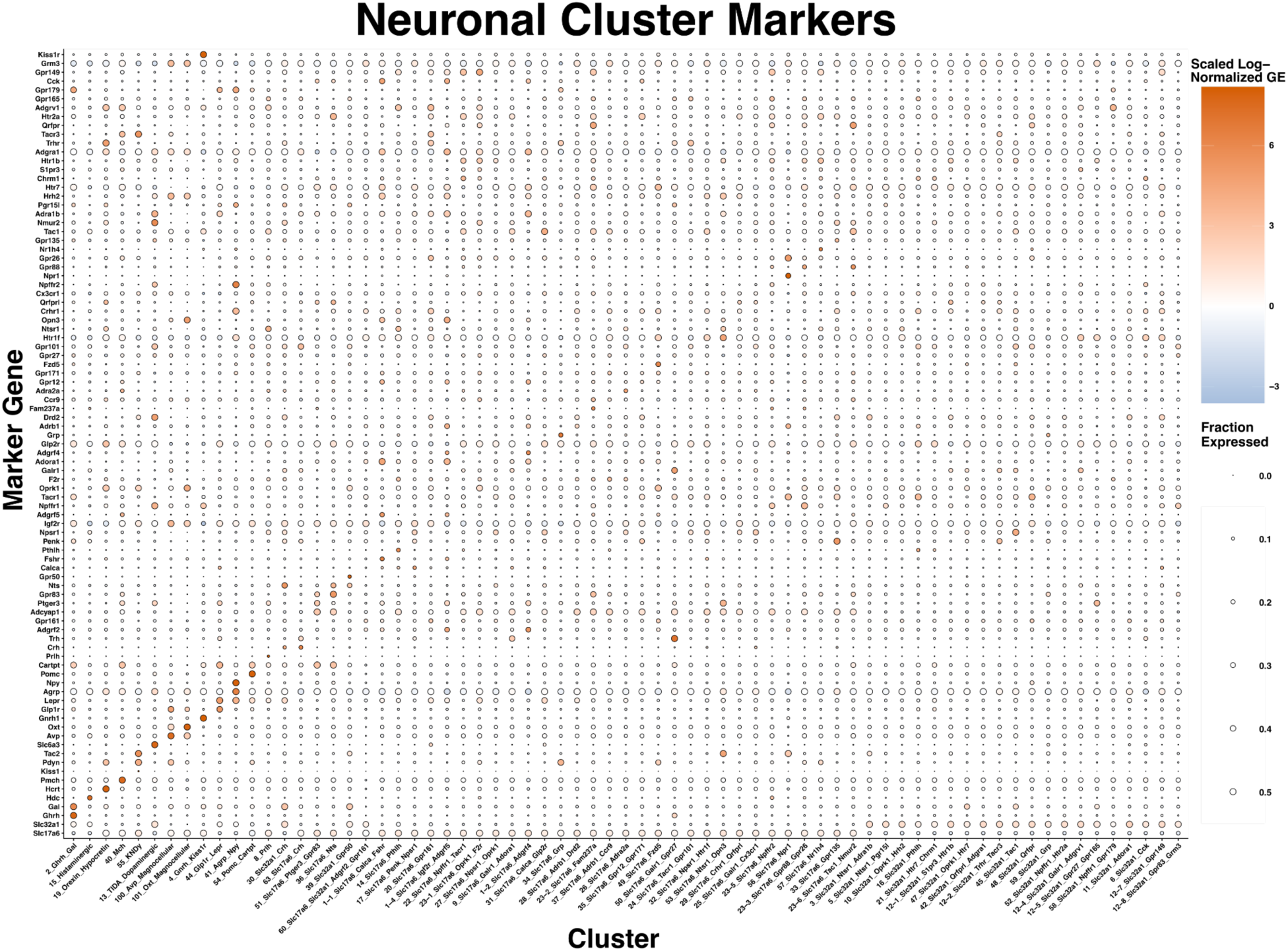
Dotplot of discriminatory neuropeptide or receptor marker gene expression across all 69 neuronal clusters. Color indicates z-scored log-normalized gene expression, and size of dot indicates fraction of cells with non-zero expression (capped at 0.5 for visualization).

**Supplementary Figure 3.**
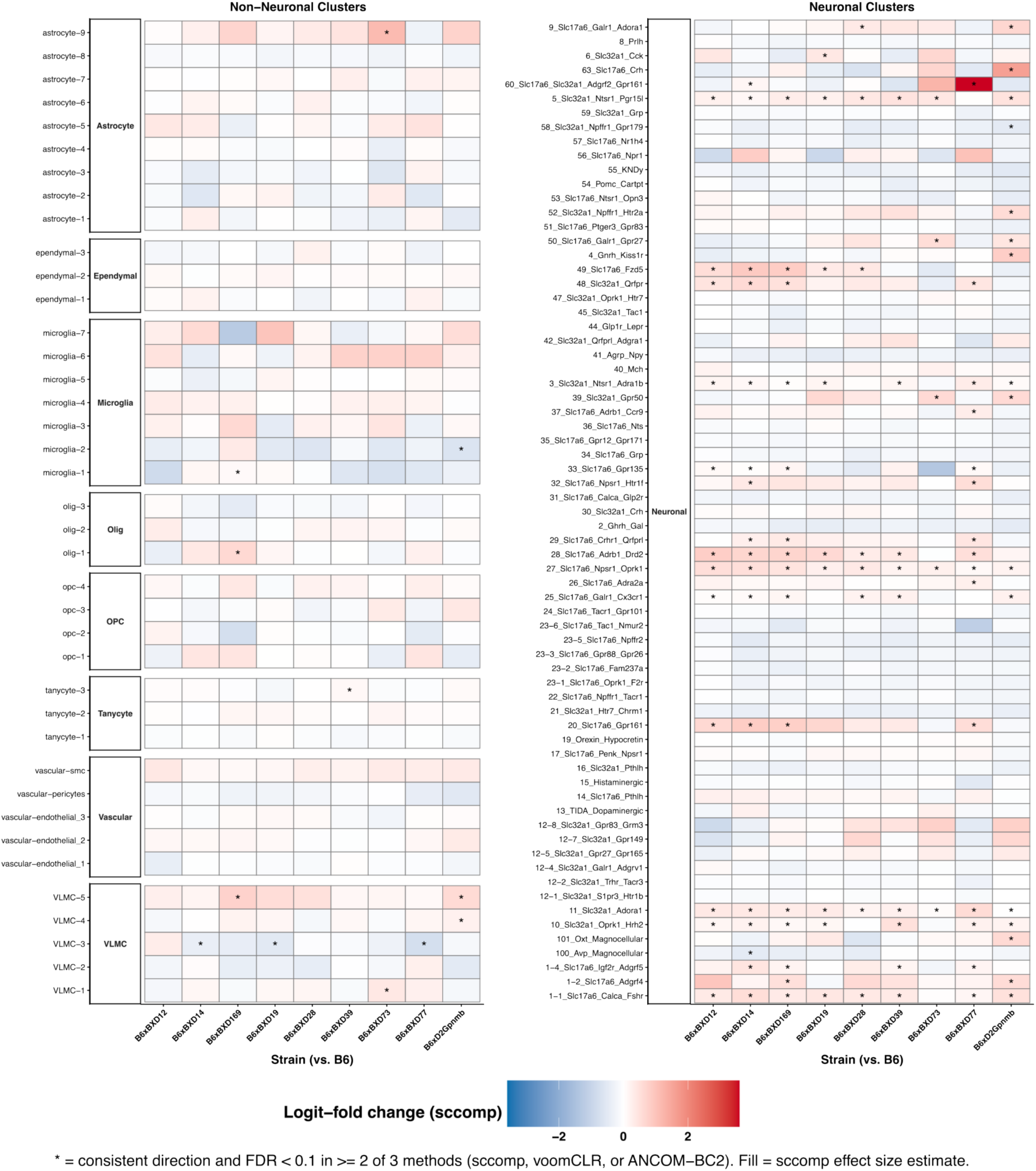
Genetic background is associated with baseline cluster-abundance composition. Color indicates log-odds fold change values and asterisk (*) indicates statistical significance in at least two differential relative abundance methods. Each box represents a comparison of relative abundance of a given cluster in B6 to another genetic background.

**Supplementary Figure 4.**
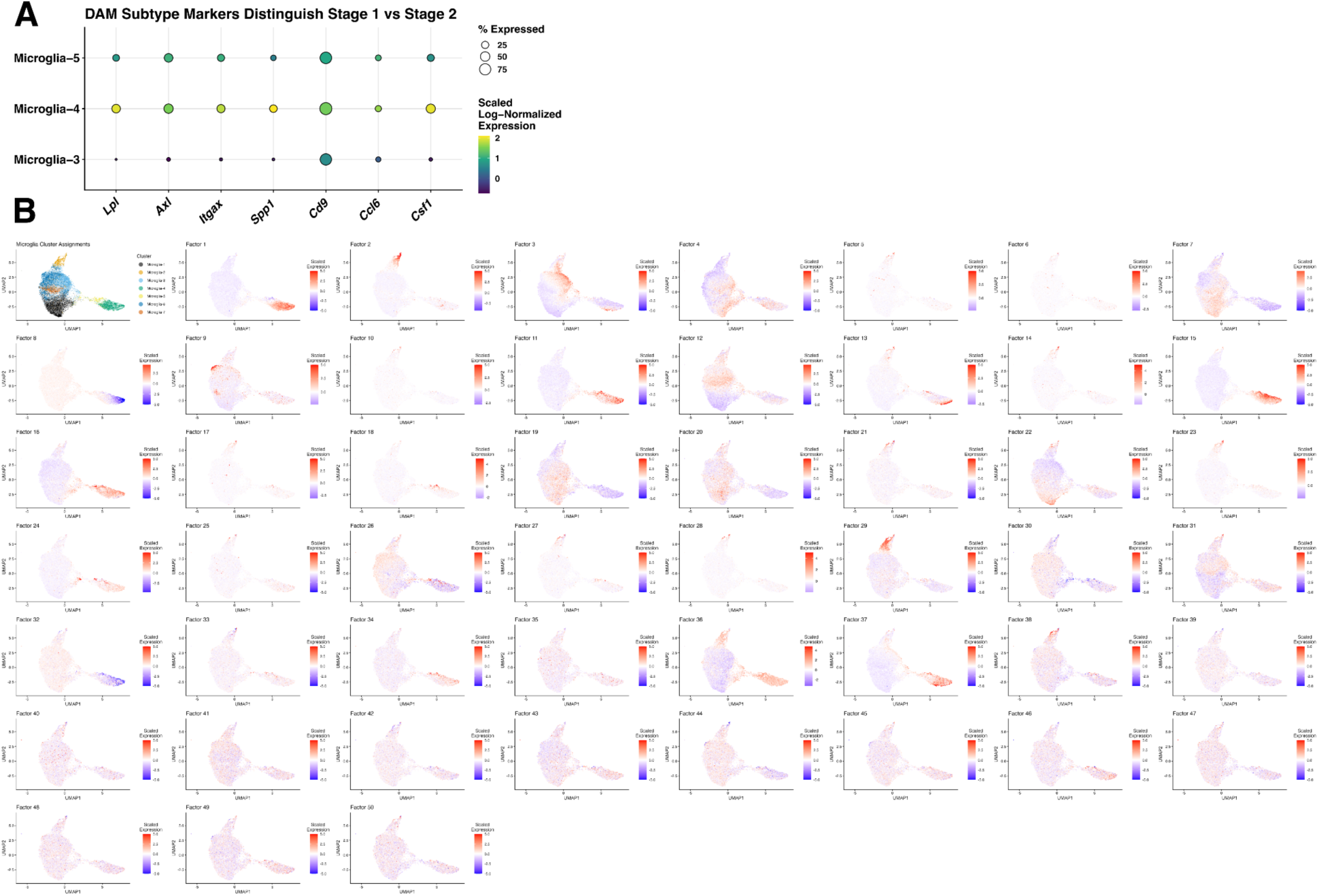
a) Stage 1 (microglia-5) and Stage 2 (microglia-4) DAM subtypes are distinguished by expression of previously defined marker genes^28^. Microglia-3, which exhibits some signatures of reactivity, is included for comparison. b) Distribution of de novo-identified factors captures cluster-spanning patterns of transcriptomic variation. The top 50 factors acquired via sciRED^32^ are included.

**Supplementary Figure 5.**
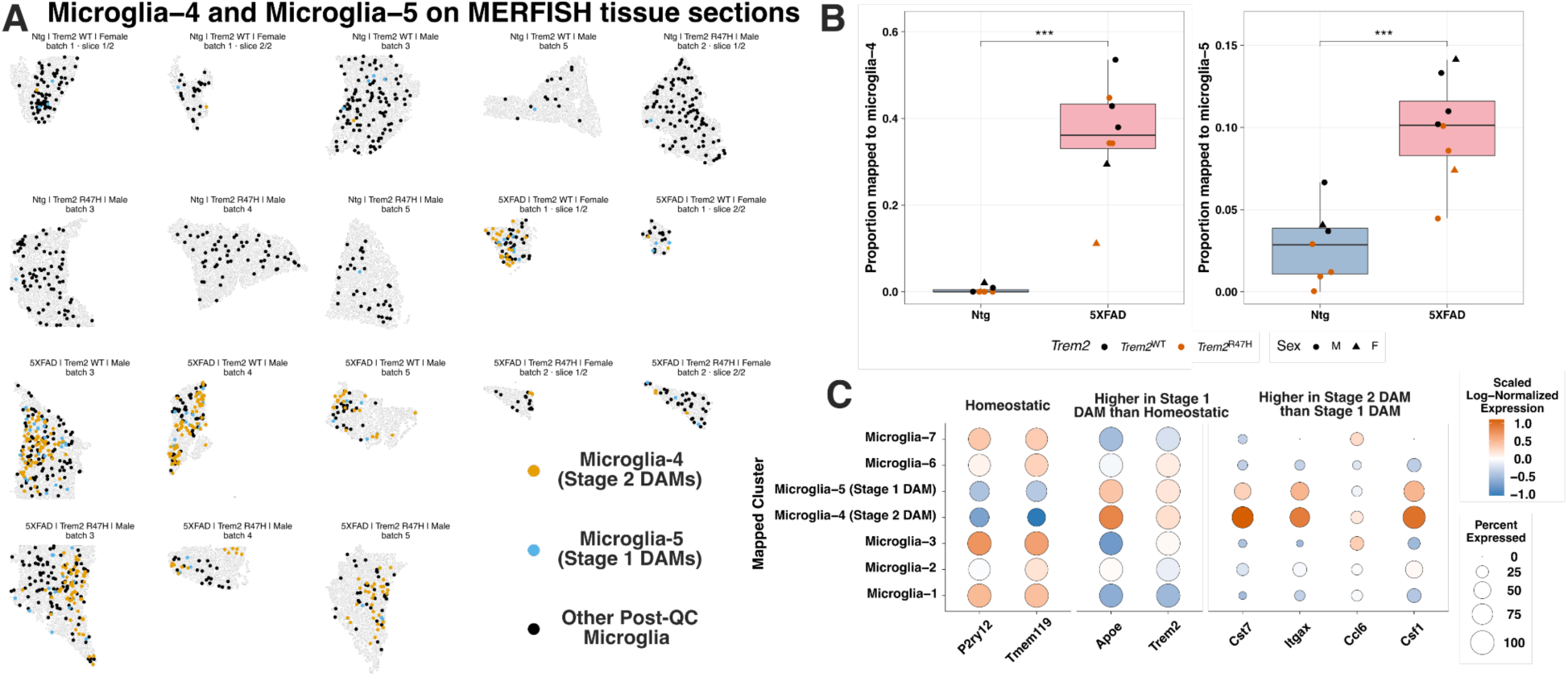
Orthogonal validation of higher Stage 1 and Stage 2 DAM relative abundance within the hypothalamus through MERFISH imaging spatial transcriptomics. a) Hypothalamic microglia profiled with MERFISH were mapped to microglial clusters, and are shown in situ alongside other hypothalamic cells. b) The 5XFAD transgene is associated with a higher relative abundance of Stage 1 and Stage 2 DAM-mapped microglia. The sccomp FDR-adjusted p-value is displayed. c) Expression of previously described^28^ marker genes that distinguish homeostatic, Stage 1 DAMs, and Stage 2 DAMs in MERFISH hypothalamic microglia mapped to snRNAseq clusters.

**Supplementary Figure 6.**
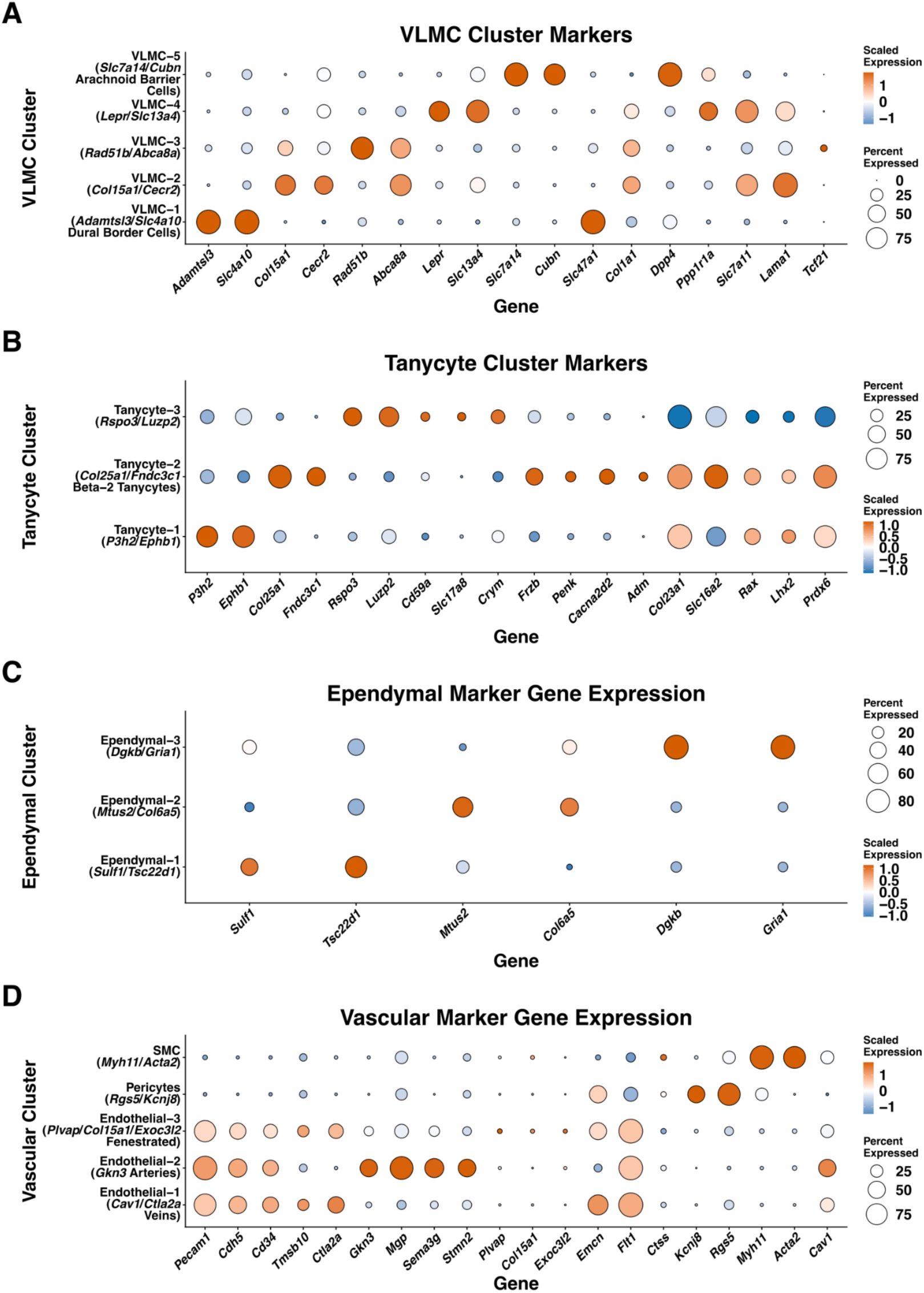
Distinct marker genes delineate clusters of a) VLMCs^102,103^; b) tanycyte cells^40^; c) ependymal cells; and, d) vascular^104,105^ cells.

**Supplementary Figure 7.**
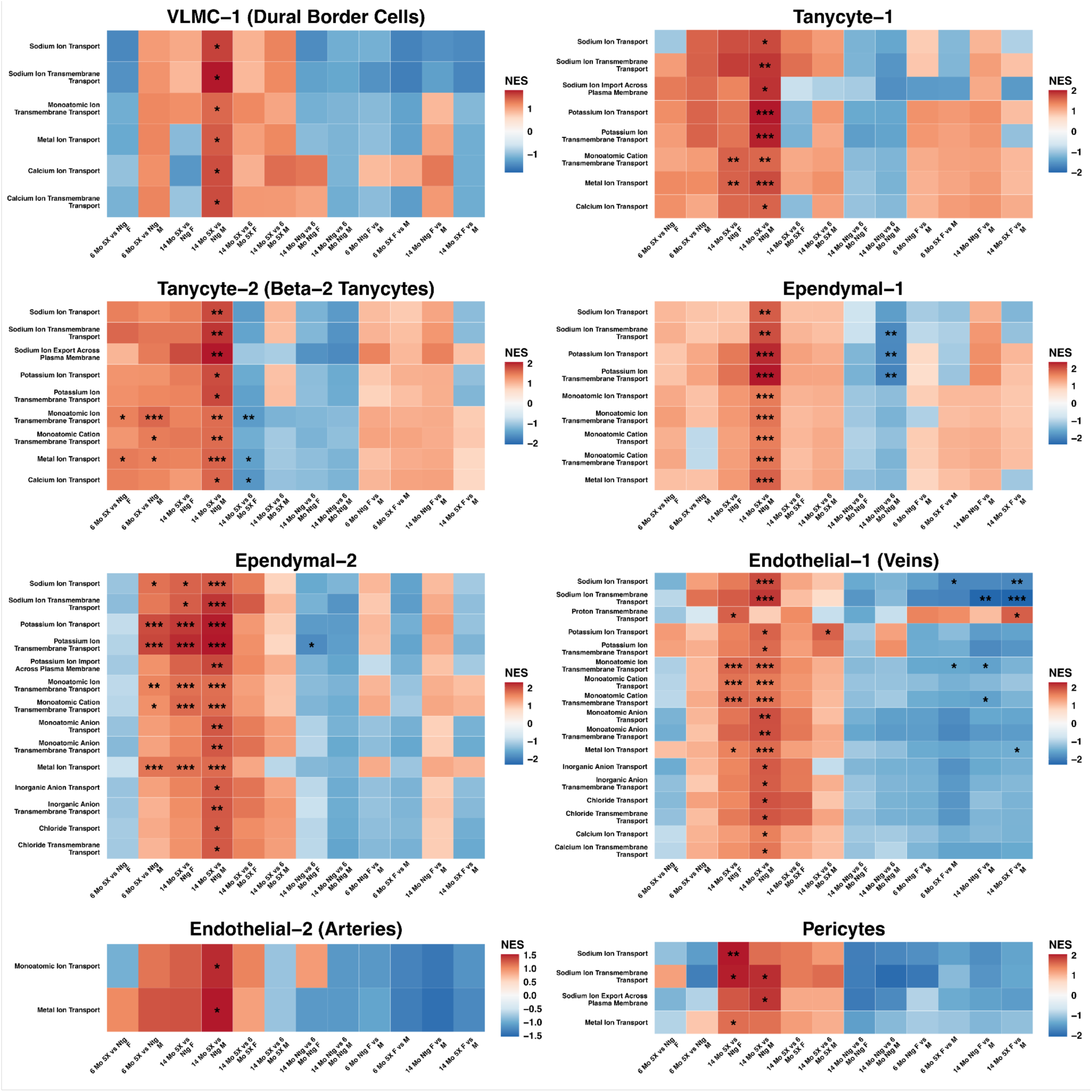
All GSEA results for barrier cell type/ion transport pathway pairs that were significantly different between 14 month 5XFAD and 14 month Ntg animals, of either sex.

**Supplementary Figure 8.**
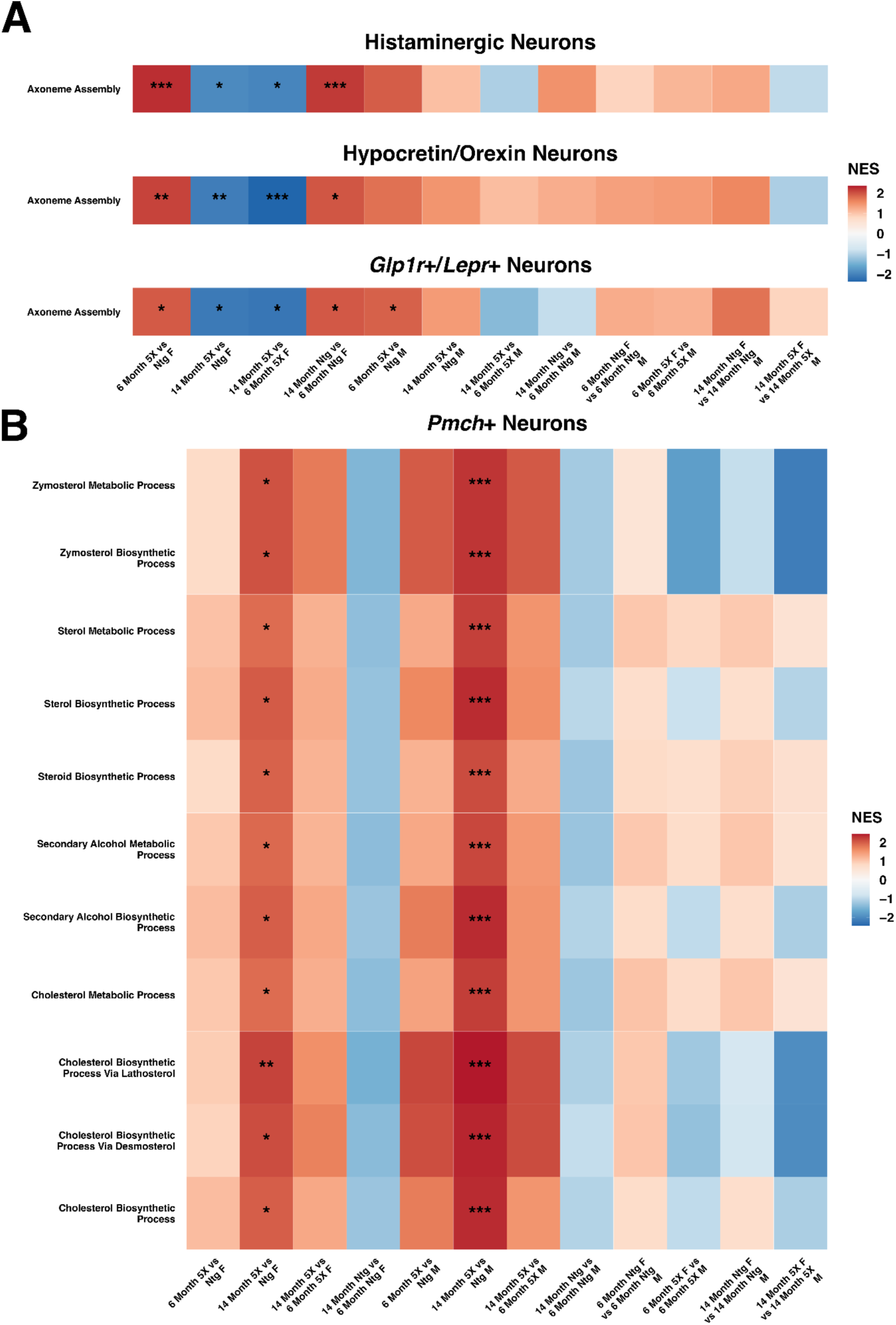
All pathway-level comparisons for highlighted a) neuronal cilia and b) Pmch+ sterol associations in Fig. 6.

**Supplementary Figure 9.**
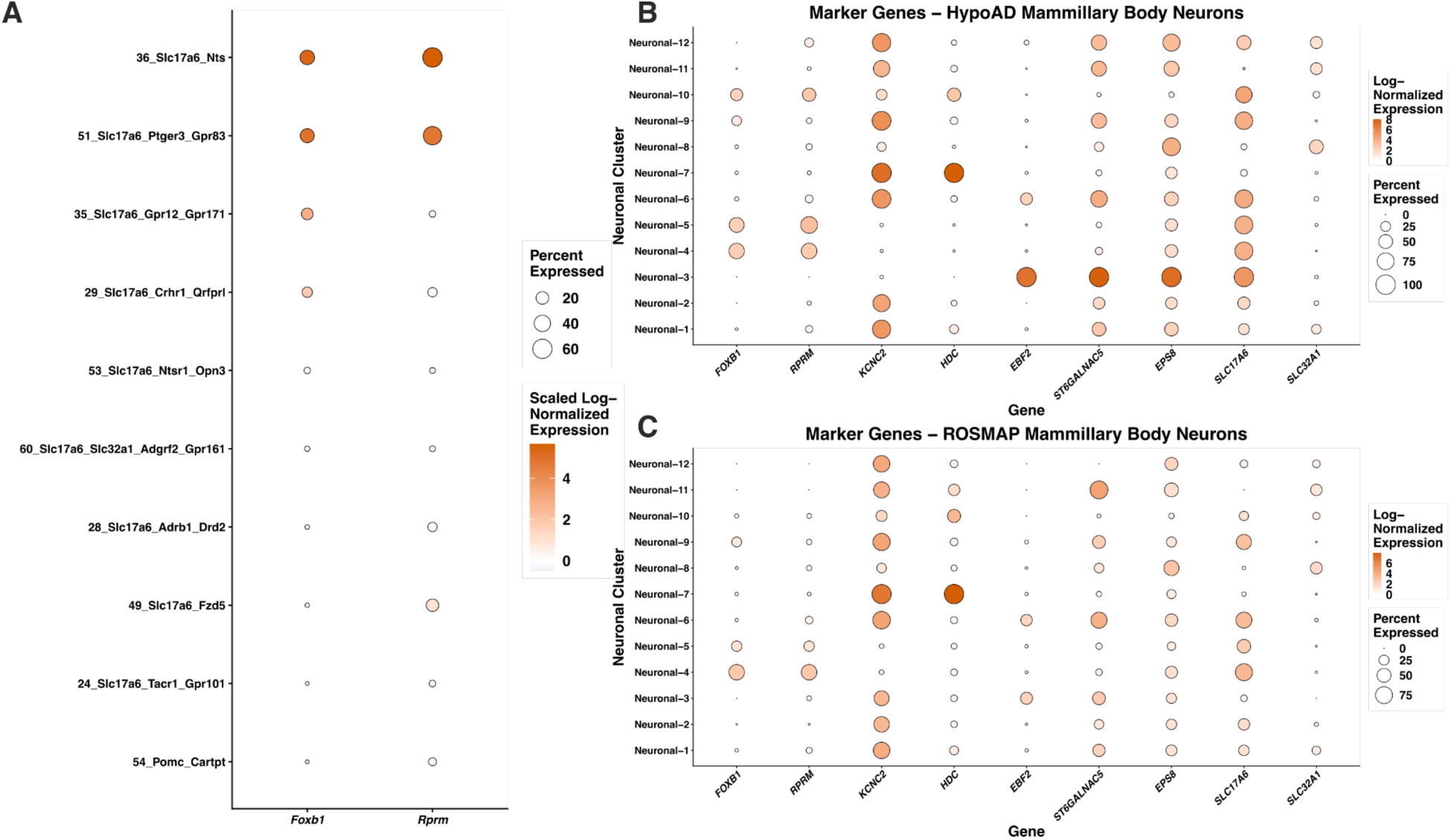
Expression of a) the medial mammillary body marker genes Foxb1 and Rprm in mice (showing the 10 clusters with the highest expression of Foxb1) and b,c) several cell type distinguishing marker genes of the mammillary bodies in the hypoAD and ROSMAP datasets.

